# Genomic Flexibility Through Extrachromosomal Amplifications: A *Leishmania* Survival Strategy

**DOI:** 10.64898/2025.12.13.692850

**Authors:** Atia Amin, Ana Victoria Ibarra-Meneses, Mathieu Blanchette, Christopher Fernandez-Prada, David Langlais

**Affiliations:** Department of Human Genetics, Faculty of Medicine and Health Sciences, Victor Phillip Dahdaleh Institute of Genomic Medicine, McGill University, Montréal, QC, Canada; Département de Pathologie et Microbiologie, Faculté de Médecine Vétérinaire, Université de Montréal, Saint-Hyacinthe, Québec, Canada; School of Computer Science, McGill University, Montreal, QC, Canada; Department of Microbiology and Immunology, McGill University, Montreal, QC, Canada

**Keywords:** Extrachromosomal amplicons, *mrpA* locus, Antimonial resistance, *Leishmania* sp., recombination mechanism

## Abstract

*Leishmania* parasites modulate gene copy number through extrachromosomal DNA (ecDNA) amplification, enabling adaptation to environmental stress. Under drug pressure, both linear and circular ecDNA amplifications (amplicons) carrying resistance genes emerge. However, how these ecDNA structures form, diversify, and coexist remains poorly understood. Here, using experimental evolution and Oxford Nanopore long-read sequencing, we show that a single clonal population of drug resistant *Leishmania* produces a variety of linear and circular amplicons. As antimonial pressure increases, linear amplicons transition into circular forms, with high-stress conditions favoring circular amplicons carrying at least two copies of the resistance gene. Using the Nanopore long reads, we map recombination events driving linear and circular amplicon formation. Our model suggests that gene duplication in the amplicons originates from inter-chromatid homologous recombination, leading to an intermediate intra-chromosomal duplication, followed by a second homologous recombination event. Additionally, different *Leishmania* species exhibited distinct biases toward linear or circular amplification under identical drug conditions, suggesting species-specific adaptive strategies. Together, these findings define recombination-driven ecDNA dynamics as a central axis of genomic plasticity in *Leishmania* and underscore the potential for targeting ecDNA in therapeutic and diagnostic strategies against *Leishmania* and related pathogens.

## Introduction

Gene expression in eukaryotes is modulated through a variety of mechanisms, including variation in gene copy number (CNV), transcriptional regulation, and post-transcriptional modifications. When one mode of regulation is limited, others must compensate to survive. *Leishmania*, an early-diverging kinetoplastid protozoan parasite and the causative agent of leishmaniasis, lacks conventional gene-specific transcriptional control due to the absence of defined promoters and typical RNA polymerase II regulation (Campbell, Thomas, and Sturm 2003; Clayton 2019). Instead, gene expression in *Leishmania* is predominantly controlled post-transcriptionally (Karamysheva, Gutierrez Guarnizo, and Karamyshev 2020) and through modulation of gene dosage, which is achieved by ploidy alteration and copy number variations (CNVs) (Iantorno et al. 2017; Ubeda et al. 2008).

One prominent mechanism of CNVs involves the generation of extrachromosomal DNA (ecDNA) amplifications (amplicons); circular or linear DNA molecules that can replicate and segregate independently of chromosomes (Hull and Houseley 2020; Ubeda et al. 2014). While ecDNAs have been described over a half a century ago, their biological significance has gained renewed attention in the past decade with several studies showing their implication during genomic instability, stress response, and potential as diagnostic biomarkers for cancers and other diseases (Yang et al. 2025). Functions of ecDNA have been investigated across diverse eukaryotes. For example, in fungi, ecDNA molecules can emerge as a mechanism for adaptation to nitrogen-limiting environments (Gresham et al. 2010). Recently, their role in genomic instability and stress responses has been investigated in plants (Zhuang et al. 2024; Zhang et al. 2023) and the molecular mechanisms underlying ecDNA formation have been explained in mammals (Chung et al. 2025). In cancer, ecDNAs are recognized as key drivers of oncogene amplification, tumor evolution, and drug resistance (Yang et al. 2025), as they exhibit dynamic structural plasticity, enabling rapid adaptation to selective pressures (Kim et al. 2020; Zhao et al. 2022). Recently, one study showed that structural heterogeneity of ecDNAs carrying an oncogene (*EGFR*) correlates with glioblastoma patient survival (Tang et al. 2025). Yet, despite their biological and functional relevance in diverse eukaryotes, the role of ecDNAs in parasite, mechanism of ecDNA formation, and their contribution to evolution and drug resistance remained poorly understood.

In *Leishmania*, ecDNA amplifications have been reported in association with resistance to multiple drugs. They were first described in the late 1980s, primarily in the context of methotrexate (MTX) resistance (Petrillo-Peixoto and Beverley 1988; Hightower et al. 1988; Garvey and Santi 1986; White et al. 1988). Seminal studies revealed both circular and linear amplicons carrying amplified copies of the ornithine decarboxylase gene (*ODC)* in difluoromethylornithine (DFMO) resistant *L. donovani* (Hanson et al. 1992), co-existence of both circular and linear amplicons carrying pteridine reductase gene (*PTR1*) in MTX resistant *L. tropica* (Olmo et al. 1995), and identified short homologous sequences as potential drivers of their formation via recombination (Navarro et al. 1994; Grondin, Roy, and Ouellette 1996). These early insights provided the first mechanistic clues linking extrachromosomal amplification to drug resistance in protozoan parasites. Subsequent work expanded this model to include additional resistance-associated genes such as *DHFR-TS* in MTX resistant *L. infantum* and *L. major* and showed that amplicons could persist independently of chromosomes in antimony resistant *L. infantum* with the amplification of the pentamidine resistance protein 1 gene, known as *mrpA* (Ubeda et al. 2008; Leprohon et al. 2009).

However, while these pioneering studies established the foundational principles of ecDNA biology in *Leishmania*, they were limited by the molecular tools available at the time. Most studies relied on Southern blotting, pulsed-field gel electrophoresis, DNA microarray which constrained the ability to resolve the full structure, diversity, and dynamics of amplicons. Furthermore, the broader genomic context of amplicon formation including the role of recombination mechanisms, the structural transitions between linear and circular forms, and the longitudinal evolution of amplicons under selective pressure remained largely unexplored. As a result, critical questions about how *Leishmania* dynamically remodels its genome through ecDNA amplification during adaptation and resistance acquisition have persisted for decades.

A resurgence of interest in *Leishmania* ecDNA biology followed the advent of high-throughput sequencing and the application of experimental evolution strategies. One study demonstrated that the parasite genome is stochastically and constitutively rearranged at repeated sequences, which serve as hotspots for the formation of both circular and linear amplicons under selective pressure (Ubeda et al. 2014). These amplicons can arise via RAD51-dependent homologous recombination between direct repeats, or by alternative mechanisms involving inverted repeats. Importantly, bioinformatic screens revealed thousands of repeat alignment groups across *Leishmania* species, suggesting that most of the genome has the potential to give rise to amplicons (Ubeda et al. 2014). However, despite these significant advances, several important questions remain unanswered.

Elucidating the genomic architecture of ecDNA requires accurate breakpoint detection, but short-read sequencing is inherently limited in this ability (Zhu et al. 2024). Therefore, using only short-read sequencing to study the complex structures of ecDNA might misclassify amplicon forms (Tang et al. 2025; Hung et al. 2021). To address this limitation, recent ecDNA studies have used long-read sequencing, which enables accurate detection of complex structural rearrangements (Hung et al. 2021; Tang et al. 2025). Since prior studies on *Leishmania* ecDNA relied on short-read sequencing, they were unable to capture the true diversity and dynamics of amplicons, particularly due to the large repeated sequences and tandem duplications. Therefore, whether a single gene locus indeed generate a diverse pool of amplicons, and the circular and linear amplicons can co-exist within a specific clonal population of parasites remains unclear.

Our study addresses these critical gaps by focusing on the *mrpA* locus, a prototypical ecDNA hotspot linked to antimonial (Sb) resistance. By leveraging Oxford Nanopore long-read sequencing to monitor the emergence of amplicons across independently evolved clones of *L. infantum* and *L. major* under progressive drug stress, we capture the structural dynamics of amplicon formation with unprecedented resolution. This work provides new insights into the recombination mechanisms underlying ecDNA architecture and contributes to a broader understanding of how *Leishmania* parasites harness genome plasticity to survive in hostile environments.

## Results

### Identification of repeated sequences and new junctions for circular *mrpA* amplicons

The role of the ABC transporter gene *mrpA* in the development of Sb resistance in *Leishmania* species has been well studied (El Fadili et al. 2005; Leprohon et al. 2009). Amplification of the *mrpA* gene on chromosome 23 can occur through several mechanisms, either at the level of whole chromosome via aneuploidy and/or at the level of specific genomic region via extrachromosomal circular amplicon formation (Douanne et al. 2022; Leprohon et al. 2009). Previous studies described amplicon formation in *L. infantum* Sb resistant strain Sb2000.1 at the *mrpA* locus in chromosome 23 involving four genes, a hypothetical gene (LINF_230007600), *yip1* (LINF_230007700), *mrpA* (LINF_230007800), and *arg* (LINF_230007900) (Figure 1A) (Leprohon et al. 2009). Using DNA microarray, Leprohon *et al*. 2009 identified ∼1.4 kb repeated regions flanking the *mrpA* locus which formed a circular amplicon via homologous recombination upon Sb selection. However, the precise identity of the repeat pairs involved in these recombination events and the formation of multiple circular amplicons had not been clearly established.

**Figure 1.**
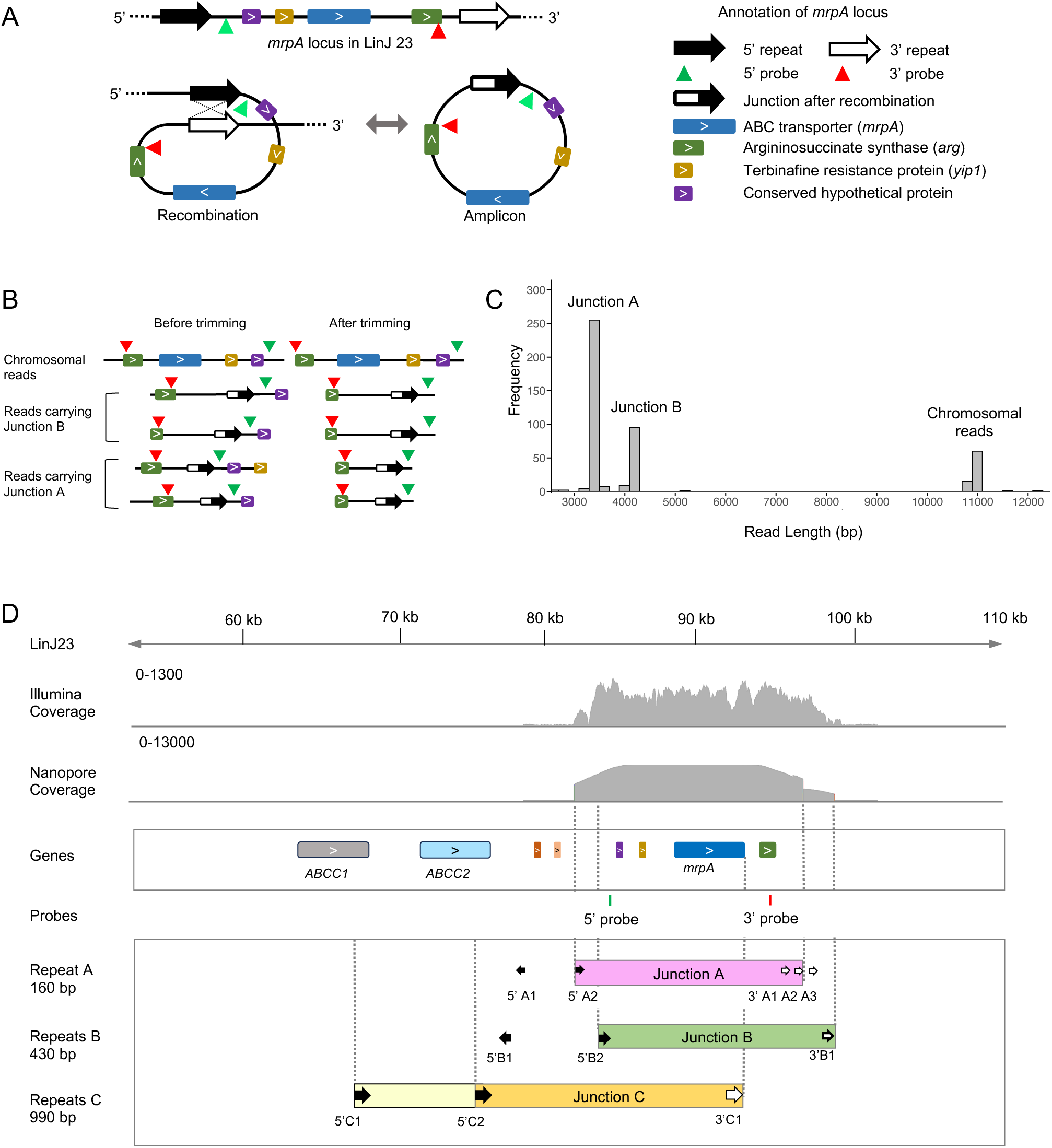
Analysis of the amplicon junctions in Sb resistant *L. infantum.* **A)** *mrpA* locus and the formation of circular amplicon through homologous recombination. **B)** A probe-based method to identify the *mrpA* amplicon junctions in an antimonial resistant strain Sb2000.1 using ONT reads **C)** Histogram showing the read length distribution of the two junctions, junction A and junction B. **D)** Analysis of the amplicon junctions in an Sb resistant strain, Sb2000.1. The ONT and Illumina read coverage tracks show the possible start and end positions, and the corresponding repeats at those positions of the two amplicons. ONT reads showed cleaner boundaries and uniform coverage compared to the Illumina coverage. Positions of repeat C were shown for reference.

The presence of a large (∼1.4 kb) repeat pairs flanking the *mrpA* locus, exhibiting >95% sequence identity (Leprohon et al. 2009), led us to hypothesize the potential formation of multiple circular amplicons at this site. We therefore performed Oxford Nanopore Technology (ONT) long read and Illumina short read sequencing to investigate this further. We designed a probe-based computational approach (Figure 1B; see Methods) to precisely capture the ONT reads corresponding to the different amplicon junctions. Histograms of the trimmed reads containing the junction sequences revealed two distinct peaks corresponding to junction A (∼3.5 kb) and junction B (∼4.2 kb), while another peak at ∼11 kb represented the reads from the chromosomal *mrpA* locus (Figure 1C). These findings provide direct evidence that multiple circular amplicons formed at the *mrpA* locus in the Sb resistant *L. infantum* laboratory strain Sb2000.1.

Upon identification of the two amplicon junctions, we wanted to precisely identify the repeats forming those junctions. Therefore, we investigated the repeat structures of the *mrpA* locus sequences in chromosome 23 in the *L. infantum* reference genome assembly (LINF GCA_900500625.2). We identified three distinct types of repeats: repeat A (160 bp), repeat B (430 bp) and repeat C (990 bp) (Figure 1D). Repeat A appears twice in opposite directions at the 5′ end, 5’A1 (LinJ23:78458-78617) and 5’A2 (LinJ23:82693-82852), and three times in the same orientation at the 3′ end, 3’A1 (LinJ23:96507-96665), 3’A2 (LinJ23: 97227-97385), and 3’A3 (LinJ23: 97947-98105). Repeat B is found twice in opposite directions at the 5′ end, 5’B1 (LinJ23: 77181-77609) and 5’B2 (LinJ23:83700-84128), and once at the 3′ end, 3’B1 (LinJ23:98952-99381). Repeat C is present three times in the same direction, twice on the 5’ end, 5’C1 (LinJ23: 67360-68368) and 5’C2 (LinJ23:75484-76471), and once at the 3’ end, 3’C1 (LinJ23:92304-93294) (Figure 1D). While repeats A and B are located in an intergenic region, repeat C is found within the *mrpA* gene itself and the two other pentamidine resistant protein genes, *ABCC1* (LINF_230007200) and *ABCC2* (LINF_230007300). Alignment of both Illumina and ONT reads showed increased coverage at the *mrpA* locus where the distinct coverage boundaries confirmed the presence of two circular amplicons, designated amplicon A and B based on the repeat types involved in the formation of the amplicon junctions (Figure 1D). In addition, alignment of the ONT read coverage boundaries with the repeat positions provided visual clues on the sets of specific repeats involved in the recombination to form the amplicon junctions (Figure 1D).

### Strategy to confirm the recombining repeats for amplicon A and amplicon B

To confirm which specific repeat pairs mediate the formation of circular amplicons A and B, we first performed multiple sequence alignment (MSA) of repeats A and B to identify sequence differences that could be leveraged to distinguish them. For repeat A, we observed distinct mismatches that differentiate the repeats based on their genomic positions and orientations: a C>A mismatch at the start and a C>T mismatch at the end distinguish the forward-facing 5′ and 3′ repeat copies (Supplementary Figure 1A). Similarly, for repeat B, we identified a characteristic single nucleotide mismatch (G>T) at the 3′ end that distinguishes the 5′ and 3′ repeats (Supplementary Figure 1B). These mismatches serve as informative markers to track repeat exchanges during recombination.

Next, to investigate the precise point of recombination and the transition from 3′ to 5′ sequence during circularization, we analyzed the ONT amplicon reads captured using the probe-based method. These reads were trimmed at the 5′ and 3′ probe boundaries and aligned alongside the corresponding reference 3’ and 5’ sequences from chromosome 23 (Supplementary Figure 2A). The resulting MSA revealed that, for amplicon A, the junction occurs at the recombination site between 3′A2 and 5′A2; for amplicon B, the transition takes place between 3′B1 and 5′B2 (Supplementary Figure 2A). Prior to the junction, the reads aligned exclusively with the 3′ reference repeat, but immediately after the junction, the alignment switched to the 5′ repeat sequence. This alignment pattern provided evidence for the recombination-driven transition in read orientation, a hallmark of circular ecDNA formation through homologous recombination.

Given that recombination involves an exchange between two repeat sequences with specific combinations of mismatches, we hypothesized that the resulting amplicon junctions should contain a combination of unique sequence features present in their parental repeats (Supplementary Figure 2B). To test this, we analyzed the junction-spanning reads for single nucleotide polymorphisms (SNPs) that could validate the recombination events. Indeed, when the amplicon reads were aligned to the reference *mrpA* locus, the junctions exhibited the predicted distinct mismatches, which appeared as SNPs in the Integrative Genomics Viewer (IGV) (Robinson et al. 2011) (Supplementary Figure 2C). Altogether, these results validate that repeat pair 5′A2 - 3′A2 mediate the formation of amplicon A, while 5′B2 - 3′B1 mediate the formation of amplicon B, both via homologous recombination.

### Sb-resistant clones generate linear amplicons carrying *ABCC1* and *ABCC2*

Following our identification of multiple circular *mrpA* amplicons in *L. infantum* strain Sb2000.1, we explored the diversity of amplicon formation across different *Leishmania* species and following stepwise *in vitro* selection under increasing Sb concentrations, P0 (no drug), 1X, 2X, and 4X of EC_50_. The EC_50_ were determined as the average of the EC_50_ of the P0 clones: 150 µM for *L. infantum* and 30 µM for *L. major*. To this end, we generated ten independent Sb-resistant clones for *L. infantum* (LINF) which primarily causes visceral leishmaniasis and seven for *L. major* (LMAJ) which typically causes cutaneous leishmaniasis (Figure 2A; see Methods). The EC_50_ was measured for each clone at each step of the selection process to assess the Sb resistance. The EC_50_ increased from 150 µM to 1200 µM in LINF clones and from 20 µM to 600 µM in LMAJ clones (Supplementary Figure 3A, B). To evaluate if the increased resistance is driven by *mrpA* amplification, we measure the expression of *mrpA* by RT-qPCR, which revealed a 4-to-240-fold increase in the LINF clones when comparing the 4X to their P0 counterparts (Supplementary Figure 3C). Of note, there was no expression of *mrpA* in the LMAJ clones at P0 (Supplementary Figure 3D). This increased expression of *mrpA* associated with resistance to Sb in *L. infantum* is suggestive of ploidy increase or formation of amplicons at or near the *mrpA* locus as observed in Sb2000.1.

**Figure 2.**
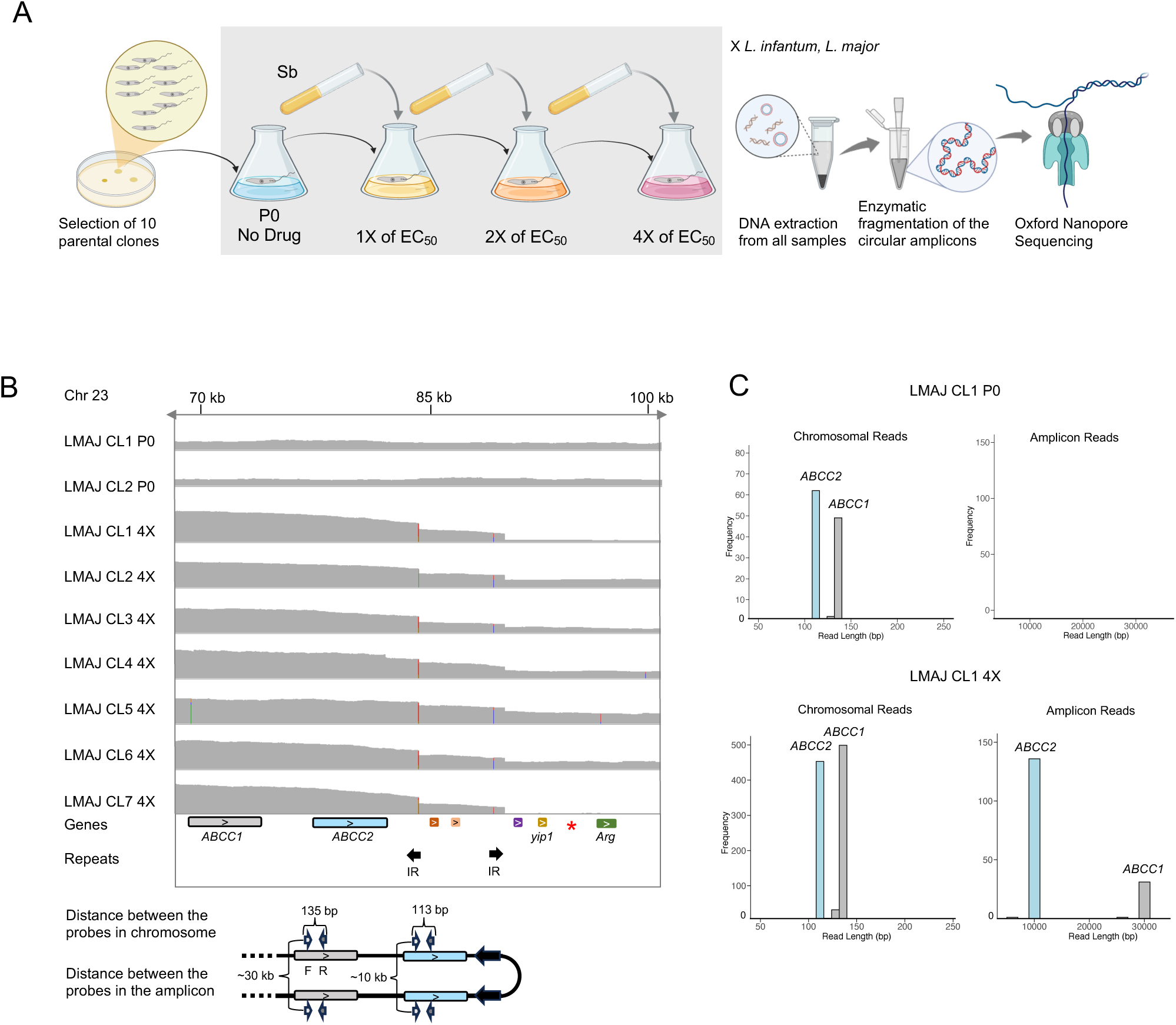
Identification of linear amplicons in Sb resistant *L. major* clones. **A**) Schematic representation of the Sb resistant clone development of two *Leishmania* species, *L. infantum* and *L. major* under four increasing Sb drug concentrations. Enzymatic fragmentation of the circular amplicons was performed during library preparation for Oxford Nanopore sequencing. **B)** *L. major* clones developed a linear amplicon carrying the *ABCC1* and *ABCC2* genes within the amplicon providing evidence for *mrpA* independent Sb resistant mechanism in *L. major*. **C)** Histograms of trimmed ONT reads show the chromosomal and amplicon reads at P0 and 4X concentrations in LMAJ CL1.

To investigate the mechanisms of resistance to Sb, we performed ONT sequencing of the different *L. infantum* and *L. major* clones. Calculating the overall sequencing coverage for each chromosome, we observed an increase in chromosome 23 in both LINF and LMAJ clones at 4X compared to their P0 counterparts (Supplementary Figure 4). A more detailed analysis of read coverage over chromosome 23 and the *mrpA* locus revealed that all LMAJ clones selected at 4X of EC_50_ Sb indeed harbored a linear amplicon (Figure 2B). Interestingly, ONT sequencing confirmed that the *mrpA* gene was completely absent from the genomes of the P0 clones picked from the *L. major* LV39 strain (Figure 2B), which agrees with our previous RT-qPCR results (Supplementary Figure 3D). Despite lacking *mrpA*, these clones exhibited a substantial increase in Sb resistance, with EC_50_ values rising up to >600 µM. Examination of the ONT read coverage, revealed linear amplicons that consistently carried two orthologs of *mrpA*: *ABCC1* and *ABCC2* (Figure 2B). The amplicon junction is flanked by inverted repeats (IRs), with SNPs arising from the exchange between these IRs clearly visualized at the amplicon boundaries on IGV (Figure 2B), further supporting the recombination-driven formation of these amplicons.

To further validate the function and structure of the linear amplicons, we first assessed the expression of *ABCC1* and *ABCC2* by RT-qPCR; all clones showed an increased mRNA expression of these ABC transporters in 4X versus unselected clones (Supplementary Figure 3E, F). We next investigated the ONT sequences, by selecting specific probe pairs within *ABCC1* and *ABCC2* with an expected distance between the respective probes of 135 bp and 113 bp (Figure 2B). In chromosomal reads (i.e. long reads extending from the ABC genes to the segment beyond 3’ IR), the observed distance between the specific probes is as expected (Figure 2C). However, in the reads derived from the amplicon, we observe distances of ∼30 kb and ∼10 kb between 2 forward probes located in *ABCC1* and *ABCC2* respectively, as expected if recombination occurred between the IR (Figure 2B, C). The parental (P0) samples showed only chromosomal reads, whereas the 4X clones displayed both chromosomal and amplicon reads, demonstrating the presence of linear amplicons encompassing *ABCC1* and *ABCC2* (Figure 2C). Identification and trimming of the ONT reads based on the start and end positions of the corresponding probes provided clear structural evidence for the identified linear amplicon, which was generated using the same set of IR in all *L. major* clones.

Although *mrpA*-independent Sb resistance mediated by *ABCC1/ABCC2* amplification had previously been reported in *L. infantum* (Douanne et al. 2020), our study is the first to demonstrate this mechanism in *L. major*. These findings highlight the evolutionary plasticity of *Leishmania* and its capacity to deploy alternative genetic strategies to achieve drug resistance in the absence of canonical resistance genes.

### Linear amplicons developed in LINF clones exhibit diversity in structure and junction formation

While we observed a uniform structure of linear amplicons in LMAJ clones, LINF clones displayed a striking degree of diversity in both linear amplicon architecture and the underlying recombination junctions. Among the ten resistant clones generated, four (CL2, CL3, CL5, and CL7) developed linear amplicons at 2X and 4X Sb concentrations, with no amplicon detected at P0 (Figure 3A). Clone 2 was already showing some reads coming from a linear amplicon at 1X (Figure 3A). At least three different rearrangement points were used during the formation of the linear amplicons at the *mrpA* locus, with one of them (IR1) used preferentially. Based on the specific inverted repeat (IR) pairs involved in their formation, we classified these linear amplicons into three distinct types: type 1, type 2, and type 3.

**Figure 3.**
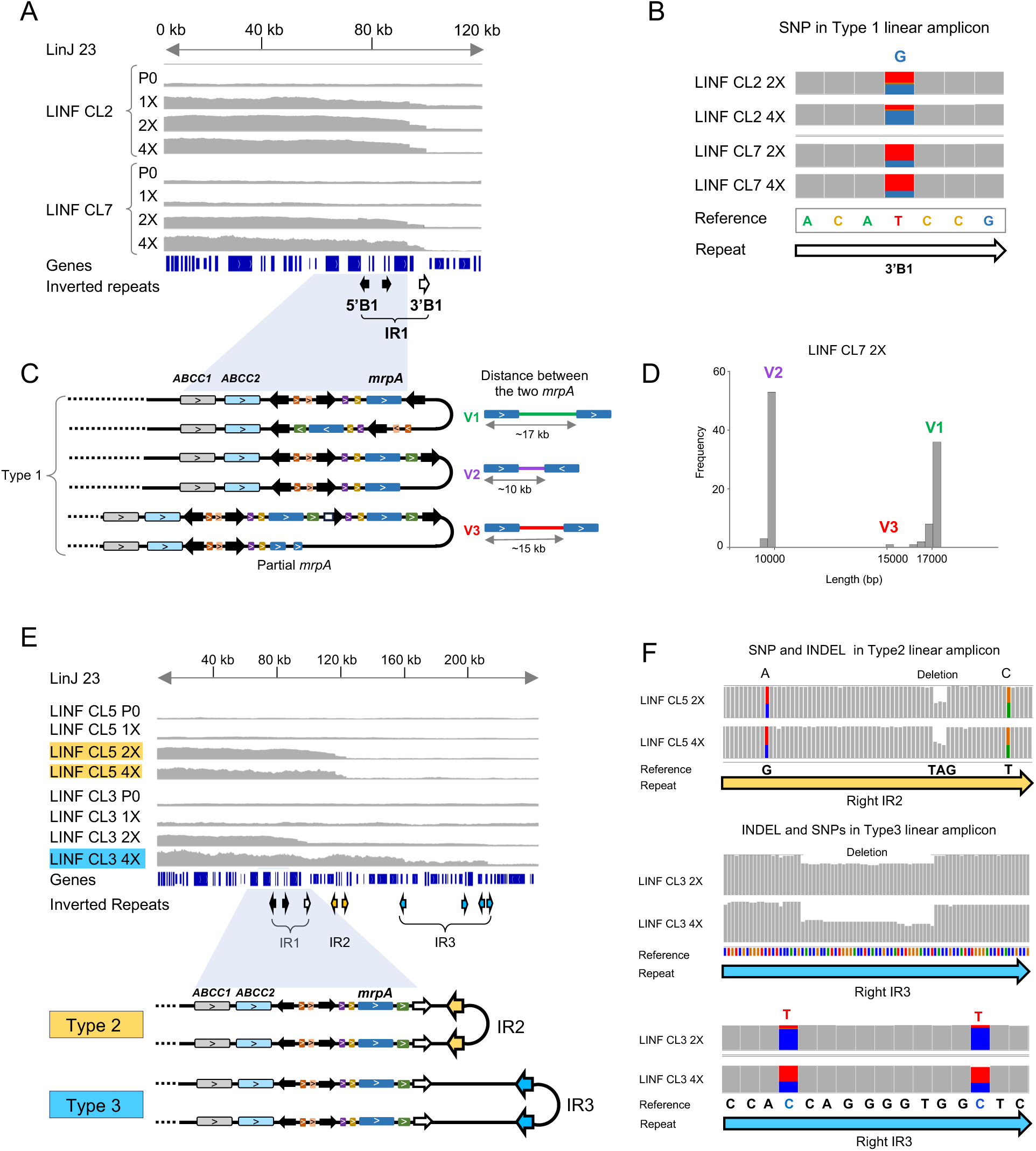
Diversity of the linear amplicons developed by LINF clones CL2, 3, 5, and 7. **A)** Increased ONT coverage indicates the presence of Type 1 linear amplicons in CL2 and CL7 at 2X and 4X. **B)** Annealing of the 3’B1 and 5’B1 inverted repeats (IR1) during the amplicon formation is evident by the T>G SNP carrying ONT reads. **C)** There are at least three variants of the Type 1 linear amplicon, V1, 2, and 3 indicated by the variation of *mrpA* locus within the amplicons. **D)** The three variants have variable distance between the two *mrpA* copies. **E)** Increased coverage indicates the presence of the linear amplicons Type 2 and 3. Inversion positions of the Type 2 and 3 linear amplicons align with the border of the corresponding inverted repeat pairs IR2 and IR3. **F)** Validation of the repeat switching between the repeat pairs, IR2 and IR3 by tracking the InDels and SNPs during the Type2 and Type3 amplicon formation respectively.

CL2 and CL7 produced only type 1 linear amplicons, which were generated through homologous recombination between the IR1, specifically 5′B1 and 3′B1 (Figure 3A). The recombination event was validated by the presence of a T>G SNP in the 3′B1 sequence, which served as a unique marker at the junction when ONT reads were aligned to the reference *mrpA* locus (Figure 3B). Further structural analysis revealed that type 1 amplicons consistently carry at least two copies of *mrpA*, however the intergenic content and orientation of the *mrpA* copies vary, giving rise to three different structural variants (Figure 3C). In version 1 (V1), the two *mrpA* copies are in the same orientation and separated by ∼17 kb; in version 2 (V2), they are in the opposite orientation and separated by ∼10 kb; and in version 3 (V3), they are arranged in the same orientations with a ∼15 kb separation (Figure 3C). Some clones contained two versions of type 1 amplicons, with CL7 containing all three, with varying frequency, while others exhibited one predominant form (Figure 3D, Supplementary Figure 5).

The remaining two clones, CL3 and CL5, also developed linear amplicons but through different recombining repeat pairs. CL3 generated a type 2 linear amplicon through recombination between IR2, while CL5 formed a type 3 amplicon via IR3 (Figure 3E). To confirm the identity of these recombining repeat pairs, we first performed multiple sequence alignments (MSA) of the IR sequences, revealing mismatches and InDels specific to each repeat pair (Supplementary Figure 6A, B). We then visualized the mutations in the ONT reads derived from the corresponding amplicons using IGV (Robinson et al. 2011), confirming the exchange of repeats through the appearance of SNPs and InDels precisely at the recombination junctions (Figure 3F).

### Formation of a double circular amplicon carrying *mrpA* in Sb-resistant LINF clones

In addition to the structural variability observed among linear amplicons, we identified significant diversity in circular amplicon formation across LINF clones. Out of ten clones, six developed circular amplicons at either 2X or 4X Sb concentrations. Notably, CL1 and CL8 transitioned from linear amplicons at 2X to circular forms at 4X, while CL6 bypassed the linear stage and formed a circular amplicon directly at 2X, which persisted through 4X without any observable linear intermediate (Figure 4A). To monitor the dynamics of amplicon formation, we tracked the allele frequency of a junction-specific SNP across increasing Sb concentrations. We observed a steady increase in the allele frequency of junction-specific SNP in CL1, 6, and 8. In CL1, for example, the allele frequency increased from 0% at P0 to 29% at 1X, 81% at 2X, and 96% at 4X—indicating that circular amplicons progressively increases in copy number, lowering the proportion of read harboring the chromosomal configuration, in a drug concentration-dependent manner (Figure 4B).

**Figure 4.**
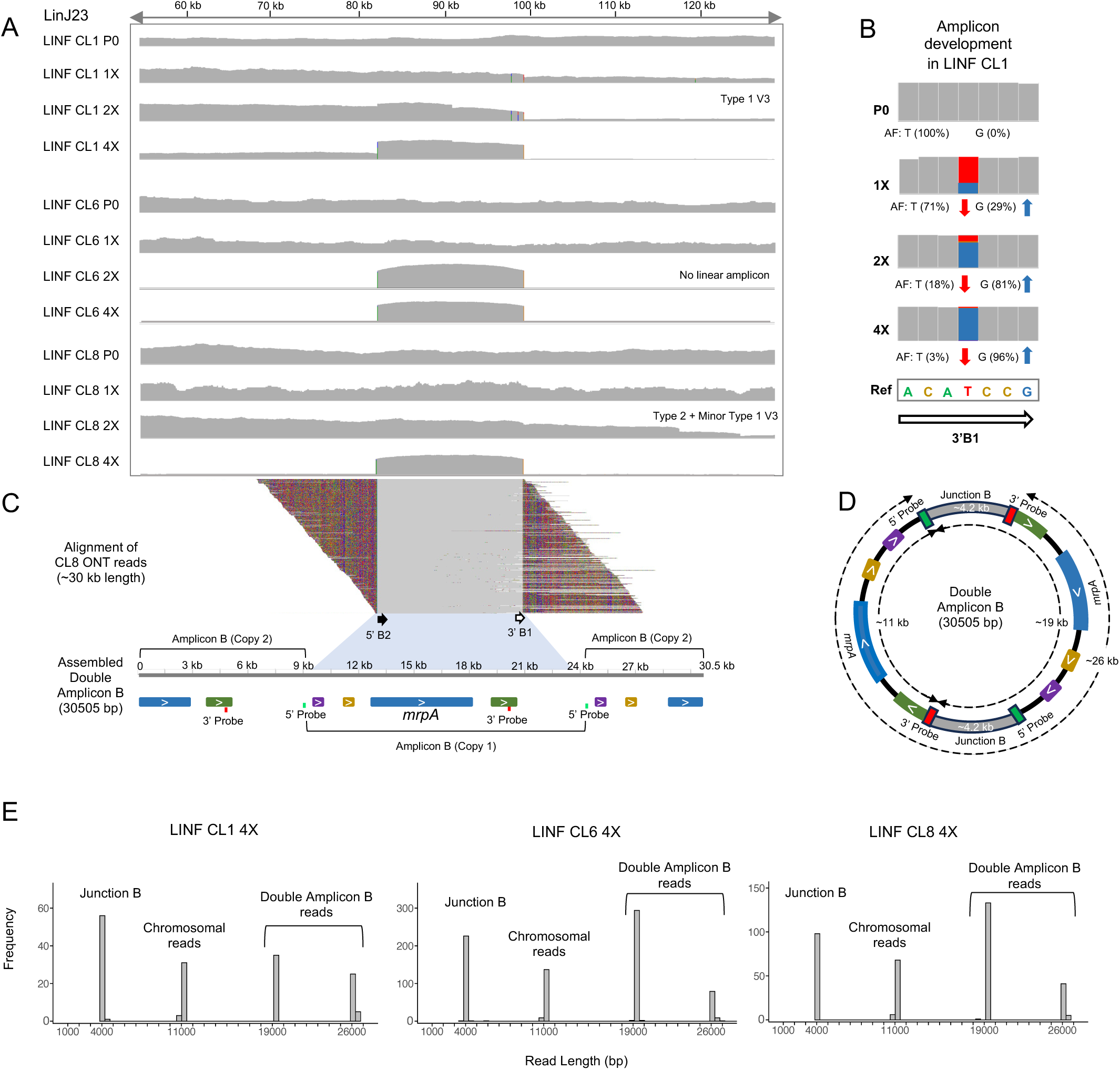
Development of double amplicon B on *L. infantum* CL1, CL6, and CL8. **A)** ONT coverage of the LINF CL1, 6, and 8 at the four Sb concentrations show the progression of amplicon development. **B)** Amplicon development can be tracked by allele frequency of a SNP at 3’B1. **C)** Assembled amplicon carries two copies of amplicon B. Alignment of individual reads carrying the double amplicon B to the *mrpA* locus in LinJ 23. The colored regions indicate the unmapped duplicated portion of amplicon reads. **D)** Distance between the two probes in the assembled double amplicon B helps track the junctions and provides evidence for the circularity of the amplicon. **E)** Junction analysis of the trimmed reads based on the probes shows the presence of junction B and the desired ∼19 kb and ∼26 kb peaks coming from the double amplicon B.

Although the circular amplicon in these clones aligned at the *mrpA* locus similarly to the previously described amplicon B and was flanked by the same repeat B sequences (5’B2, 3’B1), detailed analysis of ONT reads revealed a distinct structure. De novo assembly of the amplicon revealed a 30,505 bp contig comprising two copies of *mrpA*, two junction B sites, and the complete set of associated *mrpA* locus genes found in amplicon B (Figure 4C). This structure represents a tandem duplication of amplicon B, forming what we refer to as a “double amplicon B” (Figure 4D).

To validate the circular nature and duplication of amplicon B, we analyzed the read length distributions of ONT reads trimmed using 3′ and 5′ probes. The resulting histograms displayed four peaks, among which two at ∼19 kb and ∼26 kb were specific to the double circular configuration (Figure 4D, E). These peaks reflect the presence of reads spanning both copies of the tandem amplicon, including junctional sequences indicative of recombination events. Taken together, these findings provide evidence that CL1, CL6, and CL8 developed a novel ∼30 kb circular amplicon through duplication of amplicon B, a structure not previously described in *L. infantum* strains.

### Recombination between *mrpA* and *ABCC1*/*ABCC2* genes generates circular amplicons carrying functional *mrpA*

The final group of *L. infantum* clones, CL4, CL9, and CL10 developed a novel type of circular amplicon at 4X Sb concentration. All three clones displayed a consistent trajectory, transitioning from linear amplicons at 2X to circular amplicons at 4X, suggesting that linear-to-circular transition is a favored adaptive strategy in this strain (Figure 5A). Interestingly, the nature of the linear amplicons at 2X was varied in the three clones. Unlike the previously characterized circular amplicons A, B, and double amplicon B, this new amplicon was generated from a distinct region of the *mrpA* locus and involved the direct repeats, repeat C. Repeat C are embedded within the coding regions of *mrpA* and its paralogous transporters *ABCC1* and *ABCC2*, and their recombination gives rise to a novel junction, designated junction C. MSA of repeat C revealed two distinct regions, one with single nucleotide mismatches and InDels, and other region with 100% identity (Supplementary Figure 7A). The region with mismatches and InDels were tracked to confirm the exchange of repeated sequences during recombination.

**Figure 5.**
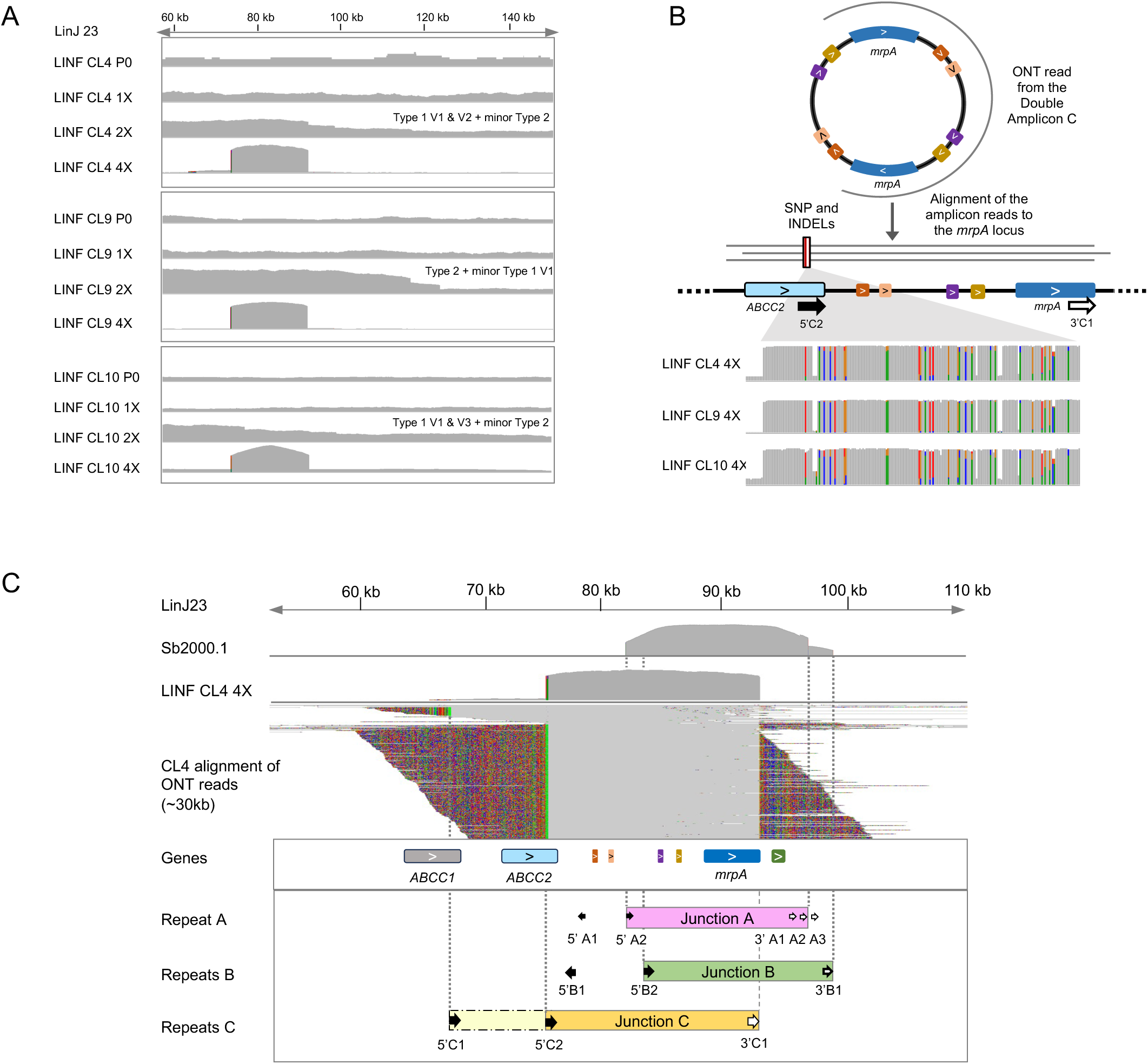
Recombination between *mrpA* and paralogous genes *ABCC1* or *ABCC2* to generate double amplicons. **A)** Progression of amplicon formation from linear to circular double amplicon C in the LINF CL4, 9, and 10 at four Sb concentrations. Linear amplicon type was indicated for 2X. **B)** Assembled double amplicon C carries two copies of identical *mrpA* as indicated by the SNPs and InDels in amplicon reads when aligned to LinJ23. **C)** ONT coverage of the lab strain Sb2000.1 shows the presence of the two other amplicons previously identified, amplicon A and B. ONT coverage of the clones show a ∼30 kb long double amplicon C which is formed through the recombination between the repeat C. Dashed line represents the second variant of the amplicon C and the corresponding repeat forming that variant.

To validate the recombination events, we performed ONT read alignments to the reference *mrpA* locus and visualized mismatches at repeat C using IGV. SNPs and InDels uniquely marking the boundaries between *mrpA*, *ABCC1*, and *ABCC2* provided evidence for sequence exchange during recombination (Figure 5B, C). Although the recombination involved coding sequences from different orthologs, the resulting amplicon C retained the full-length *mrpA* gene in all cases. This suggests a recombination mechanism that selectively preserves the functional integrity of *mrpA*, while effectively replacing portions of the *ABCC1* and *ABCC2* sequences. Supporting this, reads from amplicon C aligned across *ABCC1* and *ABCC2* in the reference genome but carried sequence features specific to *mrpA*, indicating functional replacement through repeat-mediated recombination (Figure 5B).

Interestingly, this new circular amplicon, double amplicon C, was identified in two structural variants. The more prevalent form, ∼30 kb in length, was assembled in CL4, CL9, and CL10 at 4X. It formed from the recombination between *ABCC2* (5’C2) and *mrpA* (3’C1) and thus contains two full *mrpA* copies along with four additional genes from the *mrpA* locus (Figure 5C). A shorter, less frequent ∼25 kb version was also detected in CL4, and contains a single *mrpA* gene copy, similar to amplicons A and B (Supplementary Figure 7B), likely formed from the recombination between *ABCC1* (5’C1) and *mrpA* (3’C1) (Figure 5C).

In summary, we identified five distinct circular amplicons originating from the *mrpA* locus, formed via three different recombination junctions—junctions A, B, and C (Figure 5C). Among these, amplicons A, B, and C represent single copy *mrpA* ecDNAs, while the double circular amplicons B and C contain two copies of *mrpA* within a single ecDNA molecule. This repertoire of amplicon structures highlights the extensive flexibility and redundancy of ecDNA-based gene amplification mechanisms in *Leishmania infantum*, enabling robust and adaptive resistance strategies under increasing drug pressure.

### Formation of double circular amplicons identifies a novel gene duplication mechanism

The identification of structurally diverse linear and circular amplicons arising from the *mrpA* locus, particularly those carrying duplicated *mrpA* genes prompted us to investigate the underlying mechanisms responsible for their formation. We hypothesized that the generation of double-copy circular amplicons might involve a two-step process, beginning with inter-chromatid homologous recombination that results in an intra-chromosomal locus duplication. This chromosomal duplication could act as a precursor to the formation of both linear and circular amplicons containing two copies of *mrpA* (Figure 6).

**Figure 6.**
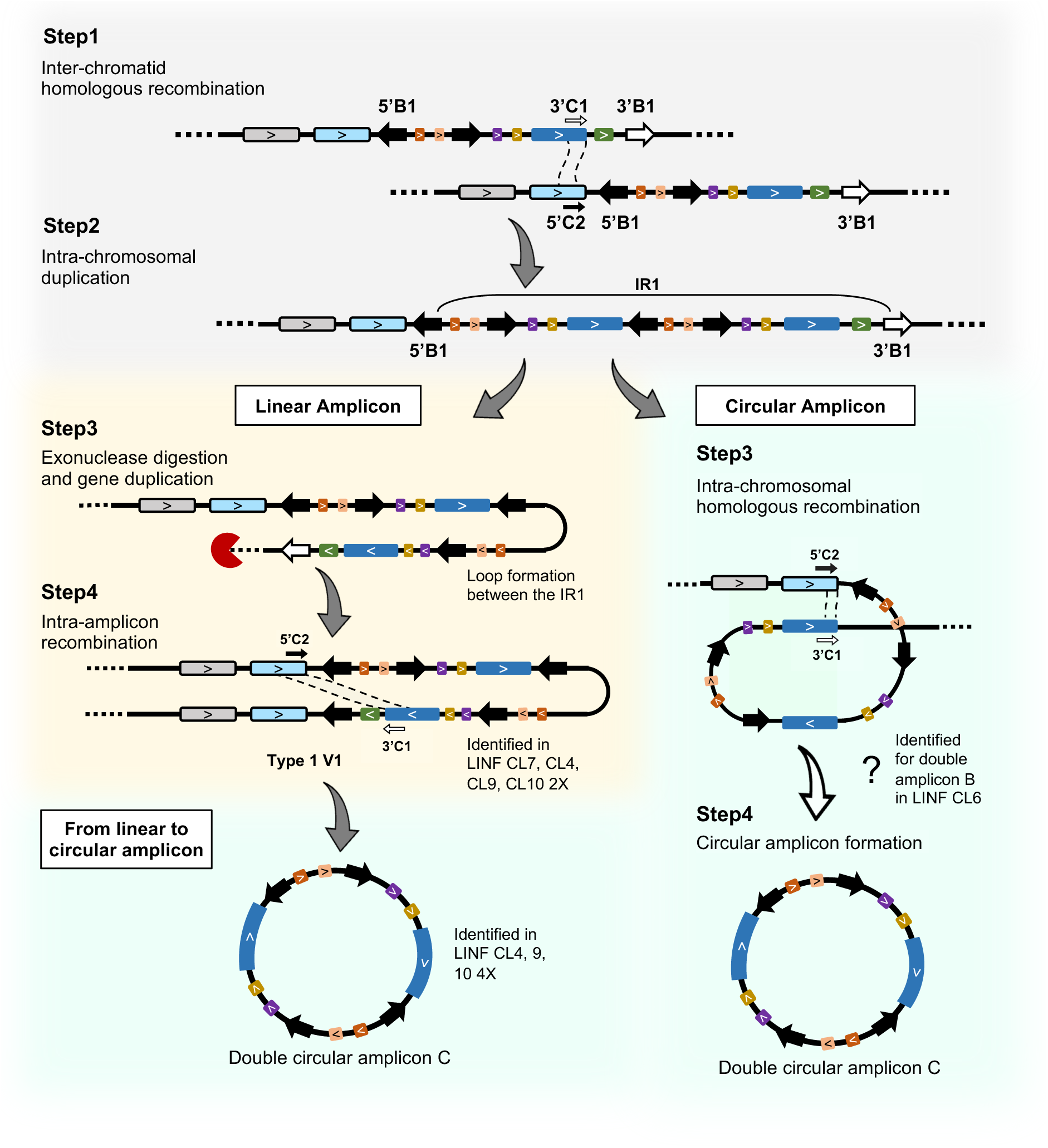
Mechanism of amplicon formation from chromosomal duplication. CL4, 9, and 10 developed linear amplicon Type1 V1 at 2X and double amplicon C at 4X indicating the progressive amplicon development from linear to circular with increasing Sb concentration. Development of linear amplicon Type1 V1 from chromosomal duplication was confirmed in CL7 2X and 4X. However, whether such chromosomal duplication can directly form double circular amplicon C could not be confirmed (indicated by a white colored arrow).

Evidence supporting this model was found in clones CL6 and CL7 at the 2X Sb concentration, where ONT reads revealed a chromosomal duplication phase. In CL7 2X, for example, ONT reads derived from the intra-chromosomal locus duplication aligned to the *mrpA* locus after the *ABCC2* gene and continued through to the 3’ telomeric end (Supplementary Figure 8A). Notably, the unmapped portion of those reads contained a duplicated *mrpA* locus, including sequences corresponding to the opposite 5’ telomeric end of chromosome 23 (Supplementary Figure 8A). The absence of this duplication phase in the other clones likely reflects the transient nature of this intermediate phase, which may exist only briefly before being resolved into extrachromosomal forms, potentially explaining why it was captured only in a subset of samples.

This chromosomal duplication phase provided mechanistic insight into the formation of the type 1 V1 linear amplicon. We propose a model where intra-chromosomal loop formation brings the inverted repeat pair 5′B1 and 3′B1 into proximity, followed by exonuclease-mediated digestion of the telomeric end. DNA replication could then proceed using the opposing telomeric sequence as a template, resulting in a duplicated linear amplicon (Figure 6). While this mechanism accounts for the formation of the type 1 V1 variant, it does not fully explain the origin of the type 1 V2 and V3 variants (Supplementary Figure 8B, Supplementary Figure 9). In the case of V2, whether the specific gene segments were lost after amplicon formation, or if the variant arose through non-homologous recombination, remained unclear (Supplementary Figure 8B).

Double circular amplicons may arise through two distinct ways. In the first, the duplication event is followed directly by loop formation and homologous recombination at direct repeat sequences, yielding a double-copy circular ecDNA without a linear intermediate. This route appears to be exemplified by clone CL6, where a chromosomal duplication was observed at 2X, and a double circular amplicon B was identified at 4X, with no intermediate linear form detected (Supplementary Figure 9).

In the second way, linear amplicons serve as transitional intermediates in the formation of double circular structures. Since linear ecDNAs or mini chromosomes already harbor the required direct repeats, they can undergo intramolecular recombination to form double circular amplicons carrying two *mrpA* copies and associated loci. This mechanism was observed in multiple clones. For example, clones CL4, CL9, and CL10 developed double circular amplicon C at 4X after forming type 1 V1 linear amplicons at 2X (Figure 6). Similarly, CL1 and CL8 transitioned from type 1 V3 linear amplicons at 2X to double circular amplicon B at 4X (Supplementary Figure 9). These observations collectively support a flexible model of ecDNA evolution in *Leishmania*, wherein recombination-driven chromosomal duplications serve as a starting point for amplicon diversification.

Overall, the structural heterogeneity of both linear and circular amplicons observed in this study, the co-existence of multiple amplicon types within the same clonal population, and the ability to form both single- and double-copy amplicons via alternative pathways underscores the remarkable plasticity of *Leishmania*’s genome remodeling machinery in response to environmental pressures such as drug exposure (Supplementary Table 1).

## Discussion

The present study provides an in-depth investigation of the structural diversity, formation mechanisms, and dynamic evolution of extrachromosomal DNA amplicons in two *Leishmania* species under drug selection, using the antimonial resistance–associated *mrpA* locus as a model. Previous studies identified ecDNA in *Leishmania* in the context of methotrexate or antimonial resistance (Grondin, Roy, and Ouellette 1996; Leprohon et al. 2009), but limitations in resolution left the locus-specific recombination and structural complexity largely unexplored. By leveraging Oxford Nanopore long-read sequencing, we uncovered multiple previously uncharacterized circular and linear amplicons originating from the *mrpA* locus, carrying single or double *mrpA* copies and formed via recombination between distinct repeat sequences.

Our findings reveal that multiple ecDNA species can coexist: co-occurring circular amplicons (e.g., amplicons A–B and C V1–V2) and linear variants (type 1 V1–V3, type 2, type 3) arise in parallel via homologous recombination between specific repeat pairs. Notably, we describe a novel double circular amplicon structure which is a duplication of either amplicons B or C—highlighting a new ecDNA architecture in protozoan parasites and supporting the hypothesis that *Leishmania* can employ recombination-mediated repeat reuse for functional gene amplification.

Our work also significantly expands the understanding of linear ecDNAs in *Leishmania*. While previous studies demonstrated the presence of linear amplicons (Ubeda et al. 2014), their architectural variation and recombination mechanism was unresolved. Here, we describe three distinct types of linear amplicons in *L. infantum* based on unique inverted repeat pairs (IR1–3), including subtype variation within type 1 amplicons, revealing the high plasticity of linear ecDNA formation. We additionally document a mechanism of linear-to-circular ecDNA evolution, as well as a model for double ecDNA formation via intra-chromosomal duplication, suggesting that ecDNA evolution can follow multi-step trajectories.

Linear-to-circular ecDNA evolution under progressive drug stress was found in *L. tropica,* and *L. tarentolae*, where the H locus first amplified as part of linear amplicons at low methotrexate (MTX) concentrations before forming circular amplicons at higher doses (Olmo et al. 1995; Grondin et al. 1998). A similar progression from linear to circular extrachromosomal DNA has been proposed in tumor cells as an intermediate step in gene amplification (Shimizu 2021), though the precise benefit of this transition remains unclear. It has been hypothesized that circular amplicons are prioritized under increased drug pressure because of its greater stability under drug pressure or the efficiency of rolling-circle replication (Beverley et al. 1984; Garvey and Santi 1986). In our work, however, we show that this linear-to-circular pathway is not universal: the LMAJ clones and four out of ten LINF clones never formed circular amplicons, even at the highest drug concentrations, while LINF CL6 produced a circular amplicon directly, bypassing a detectable linear stage.

Such heterogeneity raises the question of what governs amplicon type and variant selection. Previous studies hypothesized that culture and selection procedures could influence whether amplicons would be circular or linear (Papadopoulou, Roy, and Ouellette 1993). However, in our experiments we observed that in the identical selection procedure and culture conditions, clones develop divergent amplicon profiles. Instead, the genomic distribution of repeats appears critical: direct repeats scatter evenly across chromosomes, whereas inverted repeats cluster near sub-telomeric and telomeric regions (Black et al. 2023). In fact, the physical distance of the inverted repeats from the telomeric ends might be a factor influencing the types of linear amplicons formed (Grondin et al. 1998). In our study, 80% of the LINF clones developed at least one variant of type 1 linear amplicon, which is formed at the IR1, the nearest inverted repeat pairs to the telomeric end, indicating that telomere-proximal IRs tend to recombine more frequently.

A particularly novel finding in this study is the identification of amplicon C arising from recombination between different paralogous gene sequences (*mrpA*, *ABCC1*, and *ABCC2*). It has been reported that the repeated sequences used for the formation of extrachromosomal amplicons are generally non-coding intergenic sequences (Ubeda et al. 2008; Ubeda et al. 2014). Nevertheless, recombination between intragenic repeated regions from paralogous genes as been demonstrated in other eukaryotic cells, such as between *HXT6* and *HXT7* in Saccharomyces cerevisiae (Moller et al. 2015). In *L. tarentolae*, it has been shown that similarly to our finding, in response to arsenic exposure, a circular amplicon is generated from the recombination between *PgpA* (known as *mrpA*) and *PgpB (*known as *ABCC2)* (Ouellette et al. 1991). Interestingly, our data identified that repeats within the coding sequences of different paralogous genes can also participate in the recombination, provided that the amplicon junction preserves the gene integrity. This represents an underexplored mechanism of ecDNA creation in *Leishmania* and raises important questions about how functional integrity is preserved during recombination between homologous yet functionally divergent loci. In this case, recombination preserved *mrpA* sequence integrity within the amplicon junction, highlighting possible selective pressures for retaining resistance-conferring gene activity.

Comparative analysis across species further highlights ecDNA diversity as a versatile resistant strategy as we identified key species-specific differences in ecDNA formation. For example, while *L. major* clones developed a single type of linear amplicon, *L. infantum* clones developed a diverse type of both linear and circular amplicons. Since *L. major* lacked *mrpA* in the parental genome (P0), it rapidly acquired Sb resistance via amplification of *ABCC1*/*ABCC2*-based linear ecDNAs, indicating the ability of *Leishmania* species to flexibly reroute resistance mechanisms via alternate transporters. This type of *mrpA* independent Sb resistance mechanism was previously explained for *L. infantum mrpA* null mutant, where linear amplicons carrying *ABCC1* and *ABCC2* worked as an alternative resistance strategy (Douanne et al. 2020). Identification of similar rerouting resistance mechanism in *L. major* highlights the multifaceted nature of Sb resistance in *Leishmania* species and provides new targets for therapeutic intervention.

Overall, our results showed that *Leishmania* parasites increase drug resistance through the formation of diverse amplicons, and by expanding its genomic plasticity beyond its nuclear chromosomes. Our results support that ecDNAs in *Leishmania* are highly dynamic structures that evolve under selection pressure through diverse, repeat-mediated recombination pathways. Unlike many eukaryotes where ecDNA is associated with genomic instability and disease, in *Leishmania*, ecDNA appears to be an adaptive feature of its survival strategy, compensating for its limited transcriptional control and reliance on post-transcriptional and gene dosage-based regulation.

*Leishmania*’s remarkable genome plasticity poses a major challenge for developing effective antileishmanial drugs. By unraveling how ecDNA drives this plasticity, we can uncover key targetable dependencies for the survival mechanism in this parasite that may serve as novel therapeutic targets. Furthermore, the high-resolution amplicon structure and the multi-step recombination model made possible by long-read sequencing will accelerate biomarker discovery and guide rational drug design in *Leishmania*. Finally, these insights may extend to other rapidly evolving eukaryotes—such as cancer cells or fungal pathogens—that similarly exploit genomic instability to adapt under drug pressure.

## Materials and methods

### Sb resistant clone development of *Leishmania* species

Two strains of *L. infantum,* a wild-type strain LdiWT and an *in vitro* generated antimony resistant strain Sb2000.1 were used to screen for the presence of amplicons. Ten independent clones (P0) of *L. infantum* MHOM/MA/67/ITMAP-263 (LdiWT) and seven *L. major* LV39 promastigotes were picked form solid cultures and resistance to trivalent antimony (Sb^III^) (Potassium antimonyl tartate trihydrate, Sigma) was induced through stepwise exposure to increasing drug pressure (1X, 2X, and 4X the initial EC_50_). To assess drug sensitivity, parasites were incubated for 72 h at 25 °C in the presence of a range of Sb^III^ concentrations to calculate EC_50_ values. Parasite growth was quantified by measuring absorbance at 600 nm (A_600_) using a Cytation 5 multimode reader (BioTek, USA). Dose-response curves were generated, and EC_50_ values were calculated by nonlinear regression using GraphPad Prism 9.0 software (GraphPad Software, La Jolla, CA, USA). Three biologically independent cultures were used in each assay, which was carried out in triplicate. Statistical analyses were performed using unpaired, two-tailed *t*-tests, with significance defined as *P*-value < 0.05.

### Library preparation and DNA sequencing

We performed whole genome sequencing using the Oxford Nanopore Technology (ONT) for all *L. infantum* (P0, 1X, 2X, 4X) and *L. major* (P0, 4X) clones, as well as Illumina (MiSeq) sequencing for the two *L. infantum* strains, LdiWT and Sb2000.1. Prior to library preparation, the DNA was quality controlled on a Genomic DNA ScreenTape (Agilent TapeStation), and the libraries were prepared using the NxSeq AmpFREE kit (LCG) following manufacturer’s instructions. The indexed libraries were multiplexed and sequenced using Illumina NovaSeq 6000 in a paired-end 100 bp configuration. For long read ONT sequencing, the gDNA size was assessed using the FEMTO Pulse gDNA 165 kb kit (Agilent) and the libraries were prepared following Oxford Nanopore instructions with native sample barcoding. The libraries were pooled and sequenced on two MinION flowcells (R10.3 and R10.4), obtaining an average of 103,065 reads, 15X genomic coverage, and median Phred score of 23 per sample for *L. infantum* clones, and an average of 210600 reads, 52x genomic coverage, and median Phred score of 31 per sample for *L. major* clones (Supplementary Material 1). For LdiWT and Sb2000.1 strains, we obtained a genome coverage of 211x and 385x for ONT sequencing, and a genome coverage of 208x and 27x for Illumina sequencing respectively.

### Detection of amplicon junctions using a probe-based method

*Leishmania infantum* reference genome assembly (LINF GCA_900500625.2) was used for the analysis. For *Leishmania major*, the reference strain Friedlin (GCA_916722125.1) was used for chromosomal ploidy analysis. However, it was too different compared to our lab strain, as reflected by the numerous SNPs in the lab strain based on the ONT read alignment to the reference. Therefore, we assembled the chromosome 23 for the lab strain using Flye (Freire, Ladra, and Parama 2022) with the reads from one of the LMAJ P0 clones and used that for the amplicon investigation. The two unique probes, 5-prime probe (LinJ23:84213-84263) and 3-prime probe (LinJ23:95255-9531) were selected on the 5’ and 3’ sides of the *mrpA* respectively (Figure 1A, B). The probes were chosen as such so that they are present in the genome only once, the reads carrying both probes carry the junction of the amplicons in between, and if the reads are long enough to capture the chromosomal locus, the probes will also select the chromosomal reads (Figure 1B, C). A custom Python script (identifying_junction_reads.py, available on the GitHub repository) was used to screen the ONT reads for the presence of both the two probes and their reverse complements within the same read. The identified junction carrying reads were trimmed based on the starting position of one probe and the ending position of the other probe using the same Python script (Figure 1B). After the trimming, a sequence length distribution analysis was performed by identifying the length of the trimmed reads using SeqKit (Shen et al. 2016) and then creating a histogram using R script.

### Identification of the recombining repeats

The recombining repeats were identified in multiple steps. First, the *mrpA* locus in chromosome 23 of *Leishmania infantum* reference genome assembly (LINF GCA_900500625.2) was investigated at the UCSC genome browser (https://genome.ucsc.edu/) for the presence of simple tandem repeats. The identified repeated sequences at the 5’ end of the *mrpA* locus were searched throughout chromosome 23 using BLAST (Altschul et al. 1990). The 5’ repeats had hits on the 3’ end of the *mrpA* locus with at least 95% identity. The repeated sequences were compared for sequence identity using multiple sequence alignment (MSA) with ClustalW (Thompson, Gibson, and Higgins 2002) and pairwise comparison was performed using BLASTn. The identified mismatches between the repeats were tracked during the validation of the junction formation and recombination. We performed a MSA for junction A and B using the trimmed reads based on the probes starting and ending positions (Supplementary Figure 2A). Reads coming from the amplicons carry a unique combination of mismatches identified between the corresponding repeats if they indeed participated in homologous recombination and exchanged part of the sequence to form the junction (Supplementary Figure 2B, C). Moreover, the amplicon reads carrying the junctions will carry a mutation when aligned to the reference *mrpA* locus. We investigated and validated this mutation using IGV (Robinson et al. 2023) (Supplementary Figure 2C). The allele frequency of the mutations was recorded to track the frequency of amplicon formation at different drug concentrations.

### Ploidy calculation method

ONT sequence alignment was performed using minimap2 (Li 2018). The alignment output was processed using SAMtools (Li et al. 2009). Only primary alignment was kept. The final BAM and index files were generated for all samples. To estimate chromosomal ploidy, we computed the average sequencing depth across each chromosome. For this, BAM files were processed using BEDTools coverage (Quinlan and Hall 2010). Using the *genomecov* parameter and a genome index file, we obtained the mean per-base coverage per chromosome. For each sample, we calculated the median coverage across all chromosomes assuming most chromosomes remain diploid and used it as a normalization factor. Chromosome-wise ploidy was then estimated as:

Ploidy of a chromosome = (Mean coverage of the chromosome \ Median coverage of all chromosomes) * 2

Estimated ploidy values were rounded to two decimal places for clarity. The Python scripts for calculation and visualization of ploidy is available at: https://github.com/atiaamin2019/PampliconFinder.

### Identification of linear amplicons

Visualization of linear amplicons was performed in IGV (Robinson et al. 2023). Coverage increase throughout the amplicon boundary is observed which starts from the amplicon junction (the second IR) to the telomeric end. Because the two inverted repeats anneal at the junction, coverage increases symmetrically toward both telomeres. Mutations at the amplicon junction is also visualized in IGV after identifying them using MSA. For LMAJ, we used the *ABCC1* and *ABCC2* qPCR primers as the probes and for LINF we used the *mrpA* qPCR primers to identify the linear amplicon reads from the ONT read library. Since the linear amplicon reads should carry the forward primer and the reverse complement of the same forward primer sequence within the same read, we used them as probes to screen for the linear amplicon reads whereas the original forward and reverse primers were used to identify the chromosomal reads (Figure 2B). We also captured the reads with chromosomal duplication or linear amplicons with two *mrpA* in the same orientation (Figure 3C) by using the original forward and reverse *mrpA* qPCR primers as the probes which give the desired read length distribution after trimming the reads based on the probe position (Figure 3C, D). After that, a histogram of the trimmed reads was used to identify the linear amplicons reads compared to the chromosomal reads (Figure 2C).

### Assembly of circular amplicons

The assembly pipeline starts with the selection of the reads carrying the two unique probes (5’ and 3’) on both sides of the *mrpA* gene (Figure 1B). These probes carry the junctions in between after the amplicon is formed. After identifying the amplicon reads, those reads were trimmed so that they start with the 3’ probe and end with the 5’ probe. A sequence length distribution analysis was performed on the trimmed reads to distinguish the amplicon junctions. Reads that belong to each junction were separated. Next, each circular amplicon was assembled separately using Flye (Freire, Ladra, and Parama 2022). The options -*-meta* and *–plasmid* were selected which except uneven coverage and are appropriate for assembling circular genomes. An assembly graph is generated by Flye which was used as an input for the circularity test using Bandage (Ryan 2015). Amplicon genes and repeats were annotated using BLAST (Altschul et al. 1990).

### Validation of gene expression using qPCR

Total RNA was isolated from the cultures at P0, 4X using the RNeasy Mini Kit (Qiagen), following the manufacturer’s instructions, as described previously (Rosa-Teijeiro et al. 2021). The cDNA was synthesized with the iScript Reverse Transcription Supermix (Bio-Rad) and amplified in the iTaq universal SYBR Green Supermix Kit (Bio-Rad) using a CFX Opus Real-Time PCR System (Bio-Rad). Primers for targeted genes ATP-binding cassette protein *mrpA* (*LinJ*.*23*.*0290, LmjF.23.0250*) and pentamidine resistant genes *ABCC1* (*LmjF.23.0210*) *and ABCC2* (*LmjF.23.0220*) were designed using Primer Quest tool (https://www.idtdna.com/pages/tools/primerquest). The primer sequences of *mrpA* (*LinJ*.*23*.*0290*; Fw: 5′-CGCATTATGCTGTGGTTCCG-3′; Rv: 5′-GTCGTACTCGCCCATCAGAG-3′), *mrpA* (*LmjF.23.0250*; Fw: 5′-GGACATGGGCATCCTTGATAG-3′; Rv: 5′-CGCGGCCGACAGAATAATAA-3′), *ABCC1* (*LmjF.23.0210*; Fw: 5′-CTGCACCCATGCTGATGATA-3′; Rv: 5′-CCCACGGAATTTGCTGAAAC-3′), *ABBC2* (*LmjF.23.0220*; Fw: 5′-GCTTGGAGTTCGTGTAGAGTT-3′; Rv: 5′- AGACAAGCGCTATCCATTCC-3′). Gene expression levels were normalized to constitutively expressed mRNA encoding glyceraldehyde-3-phosphate dehydrogenase *GAPDH* (*LinJ.36.2480.*; Fw: 5′-GTACACGGTGGAGGCTGTG-3′; Rv: 5′-CCCTTGATGTGGCCCTCGG-3′); (*LmjF.30.2970*; Fw: 5′- GTACGCGAAGTAGTCAGCATTC-3′; Rv: 5′- ATCTGTGACCAGGGCCTTAT-3′). The relative amount of PCR products for each primer set was calculated based on the threshold cycle (Ct) value and the amplification efficiencies. Three technical and biological replicates were performed for each reaction.

## Data availability

All Oxford Nanopore long read sequencing data generated in this study have been deposited at NCBI Sequence Read Archive (SRA) (accession number available upon publication).

## Code availability

All custom Python and R scripts are available at https://github.com/atiaamin2019/PampliconFinder

## Acknowledgements

We thank the McGill Genome Center for supporting with the sequencing facilities. Atia Amin was supported by a Vanier Canada Graduate Scholarship. This work was supported by a Fonds de la Recherche Québec-Santé (FRQ-S) Chercheur-Boursier Jr 1 research allocation (#284497) to DL. DL is also supported by a FRQ-S Chercheur-Boursier Junior 2 Award (#348782) and the Canada Foundation for Innovation John R. Evans Leaders Fund (#38033). Work in the CFP-Lab was supported by a Natural Sciences and Engineering Research Council of Canada (NSERC) Discovery Grant RGPIN-2024-04103 and by the Canada Foundation for Innovation, grant numbers 37324 and 38858. CFP is the holder of the Canada Research Chair in Sustainable Integrative Parasitology (grant no. CRC-2024-00026).

## Author contributions

AA, CFP and DL designed the study. AA, AVIM, and DL performed the experiments, analyzed the data, and prepared the first draft of the manuscript. CFP, MB, and DL supervised the work. All authors have read and approved the content of this manuscript.

