## Supplementary Figures and Tables for "Genomic Flexibility Through Extrachromosomal Amplifications: A *Leishmania* Survival Strategy"

A

```
5' A2  >>  ATTCGGCATCCATCAAGCCAGCAGATCTGCGCACATGCACCGCCACAGCTCACCTGCCCC   60
5' A1  <<  ATTCGGCATTCATCAAGCCAGCAGATATGCGCACATGCACCGCCACAGCTCACCTGCCCC   60
3' A1  >>  ATTCGGCATCCATCAAGCCAGCAGATATGCGCACATGCACCGCCACAGCTCACCTGCCCC   60
3' A2  >>  ATTCGGCATCCATCAAGCCAGCAGATATGCGCACATGCACCGCCACAGCTCACCTGCCCC   60
3' A3  >>  ATTCGGCATCCATCAAGCCAGCAGATATGCGCACATGCACCGCCACAGCTCACCTGCCCC   60
           ********* **************** *********************************

5' A2  >>  TGCGACAACACCGCTACATCCCCGCTGTGTAGGGCGAGCTGTCTGTGGTCGAATCGGGCC   120
5' A1  <<  TGCGACAACACCGCTACATCCCCGCTGTGTAGGGCGAGCTGTCTGTGGTCGAATCGGGCC   120
3' A1  >>  TGCGACAACACCGCTACATCCCCGCTGTGTAGGGCGAGCTGTCTGTGGTCGAATCGGGCC   120
3' A2  >>  TGCGACAACACCGCTACATCCCCGCTGTGTAGGGCGAGCTGTCTGTGGTCGAATCGGGCC   120
3' A3  >>  TGCGACAACACCGCTACATCCCCGCTGTGTAGGGCGAGCTGTCTGTGGTCGAATCGGGCC   120
           ************************************************************

5' A2  >>  ATGCGCCTCACCGTCTCGTCCCGTCCTTCCTCGACCCTAC   160
5' A1  <<  ATGCGCCTCACCGTCTCGTCCCGTCCTTCCTCGACCCTAC   160
3' A1  >>  ATGCGCCTCACCGTCTCGTCCCGTCCTTCCTCGACTCTAC   160
3' A2  >>  ATGCGCCTCACCGTCTCGTCCCGTCCTTCCTCGACTCTAC   160
3' A3  >>  ATGCGCCTCACCGTCTCGTCCCGTCCTTCCTCGACCCTAC   160
           *********************************** ****
```

B

```
5' B1  <<  GCAGGGGGGGCGGCGGCGGACACGCGACTGGGGGCGGGGGTCCATGCGTCGCGCGATTCG   60
5' B2  >>  GCAGGGGGGGCGGCGGCGGACACGCGACTGGGGGCGGGGGTCCATGCGTCGCGCGATTCG   60
3' B1  >>  GCAGGGGGGGCGGCGGCGGACACGCGACTGGGGGCGGGGGTCCATGCGTCGCGCGATTCG   60
           ************************************************************

5' B1  <<  CGAACTCGCGCGTGGCACTGCGGGCCCGACACGGCGGCGGAGGCCCTGGCGGGGGTGGGG   120
5' B2  >>  CGAACTCGCGCGTGGCACTGCGGGCCCGACACGGCGGCGGAGGCCCTGGCGGGGGTGGGG   120
3' B1  >>  CGAACTCGCGCGTGGCACTGCGGGCCCGACACGGCGGCGGAGGCCCTGGCGGGGGTGGGG   120
           ************************************************************

5' B1  <<  ACGCCACCGTGCGAGCGGAGTAGCAGGCAGCGCACCACGTGCACTCCATCTTGTCACTGC   180
5' B2  >>  ACGCCACCGTGCGAGCGGAGTAGCAGGCAGCGCACCACGTGCACTCCATCTTGTCACTGC   180
3' B1  >>  ACGCCACCGTGCGAGCGGAGTAGCAGGCAGCGCACCACGTGCACTCCATCTTGTCACTGC   180
           ************************************************************

5' B1  <<  CAGTCACCCGTCGCTGCAGGAATGCAAAGGGCGGACAGGTGTGCGTGCGGAGCTGGGCGA   240
5' B2  >>  CAGTCACCCGTCGCTGCAGGAATGCAAAGGGCGGACAGGTGTGCGTGCGGAGCTGGGCGA   240
3' B1  >>  CAGTCACCCGTCGCTGCAGGAATGCAAAGGGCGGACAGGTGTGCGTGCGGAGCTGGGCGA   240
           ************************************************************

5' B1  <<  TGACGGTCTCCTGTCGTCTCGTGAGCTCGGCGCCGTCGTTGCTTGGCGCGGGACGGCTGC   300
5' B2  >>  TGACGGTCTCCTGTCGTCTCGTGAGCTCGGCGCCGTCGTTGCTTGGCGCGGGACGGCTGC   300
3' B1  >>  TGACGGTCTCCTGTCGTCTCGTGAGCTCGGCGCCGTCGTTGCTTGGCGCGGGACGGCTGC   300
           ************************************************************

5' B1  <<  CACCGTGTAACTGACGCGATTTGCCGTGAGCCGGTACACGCGTGATCGCGGTCGCGAGTC   360
5' B2  >>  CACCGTGTAACTGACGCGATTTGCCGTGAGCCGGTACACGCGTGATCGCGGTCGCGAGTC   360
3' B1  >>  CACCGTGTAACTGACGCGATTTGCCGTGAGCCGGTACACGCGTGATCGCGGTCGCGAGTC   360
           ************************************************************

5' B1  <<  TCTTCGGCACAGCGCCAGCCCAGACCGGTGTGCCGGCAGCAAGAGTCCACAGCCGTGCCC   420
5' B2  >>  TCTTCGGCACAGCGCCAGCCCAGACCGGTGTGCCGGCAGCAAGAGTCCACAGCCGTGCCC   420
3' B1  >>  TCTTCGGCACAGCGCCAGCCCAGACCGGTGTGCCGGCAGCAAGAGTCCACATCCGTGCCC   420
           *************************************************** ********

5' B1  <<  GACGGCAGGG   430
5' B2  >>  GACGGCAGGG   430
3' B1  >>  GACGGCAGGG   430
           **********
```

**Supplementary Figure 1.** The *mrpA* locus contains both direct and inverted repeats and multiple sequence alignment among these repeats reveals mismatches and INDELs. **A)** Repeat A is present twice at the 5' side and three times at the 3' side. **B)** Repeat B is present twice at the 5'side and once at the 3' side. Directionality of the repeats are indicated by double angle brackets.

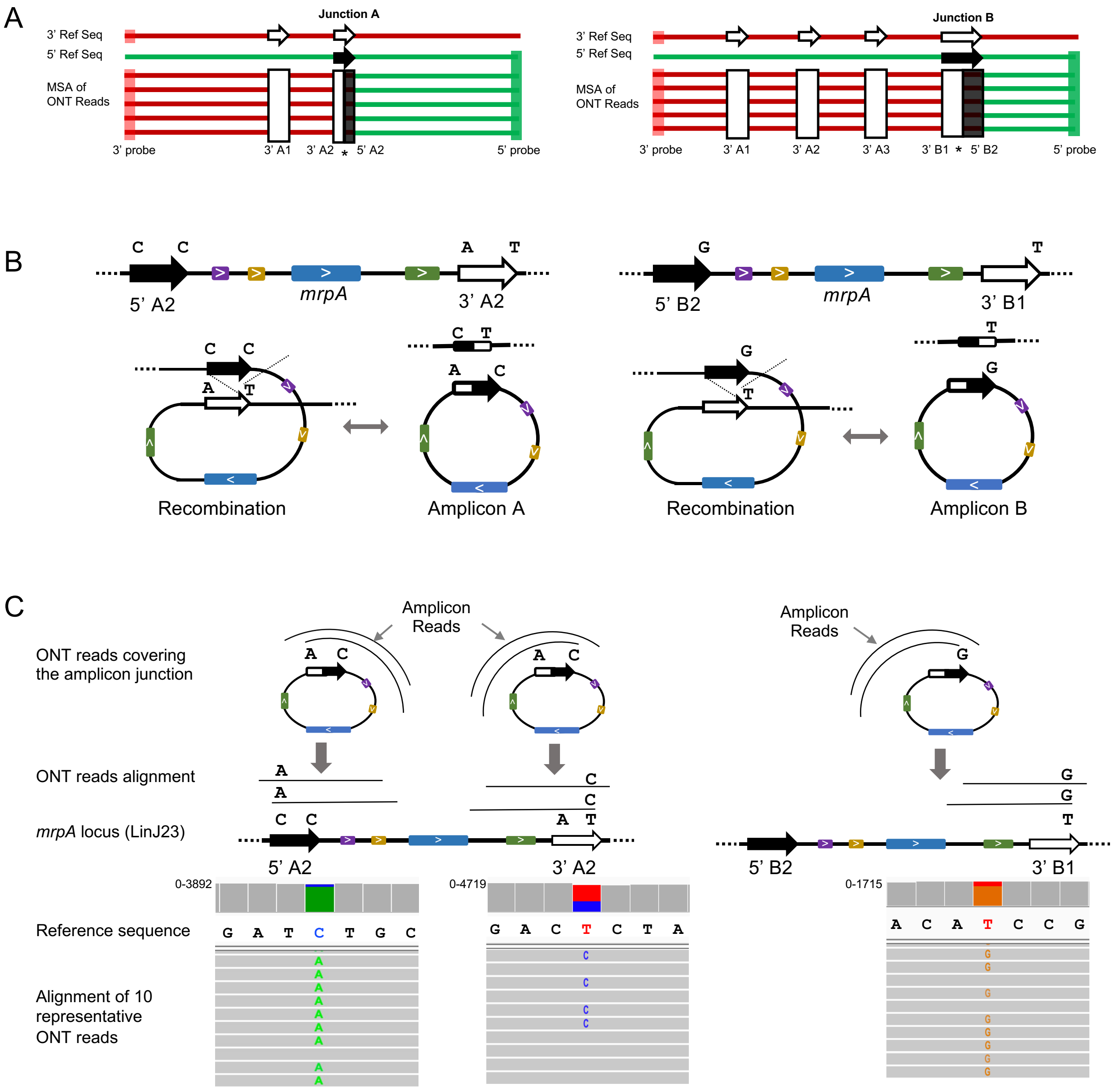


**Supplementary Figure 2.** Validation of the direct repeats participating in homologous recombination and identification of the amplicon junctions. **A)** Schematic representation of the multiple sequence alignments show the switching of the amplicon reads from the 3' to 5' side carrying the junctions in between. **B)** Schematic representation of the recombination and exchange of nucleotides between the repeats during recombination. **C)** ONT reads of the amplicons carry SNPs when aligned to the reference, confirming the exchange of the corresponding repeats during amplicon formation.

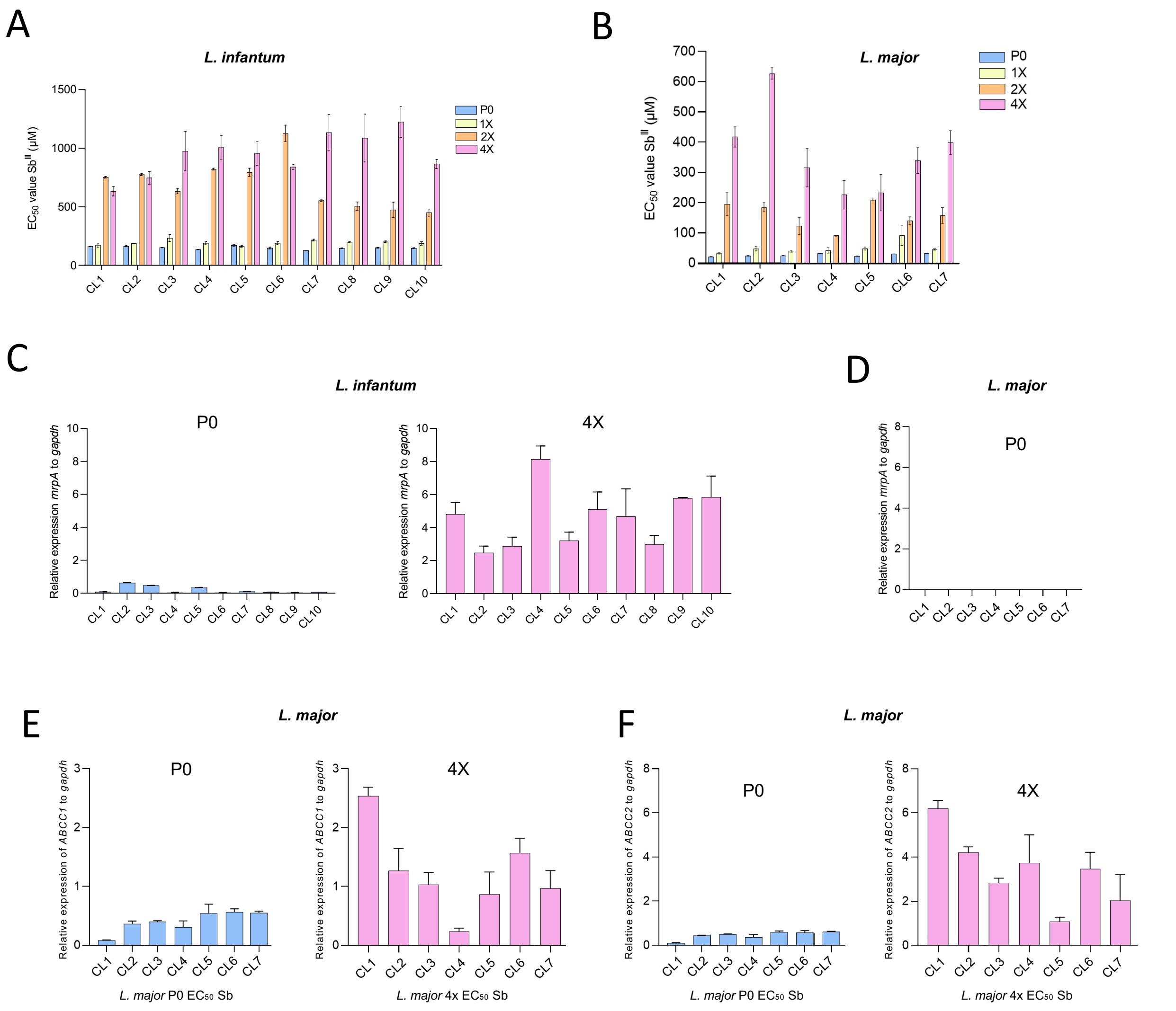


**Supplementary Figure 3**. **A, B)** EC50 of the ten *L. infantum* and seven *L. major* clones at four Sb drug concentrations. **C)** Expression of *mrpA* in the ten *L. infantum* clones at P0 and 4X Sb concentrations compared to the control gene *gapdh.* **D)** Expression of *mrpA* in the seven *L. major* clones at P0 compared to the control gene *gapdh*. **E)** Expression of *ABCC1* in the seven *L. major* clones at P0 and 4X Sb concentrations. **F)** Expression of *ABCC2* in the seven *L. major* clones at P0 and 4X Sb concentrations.

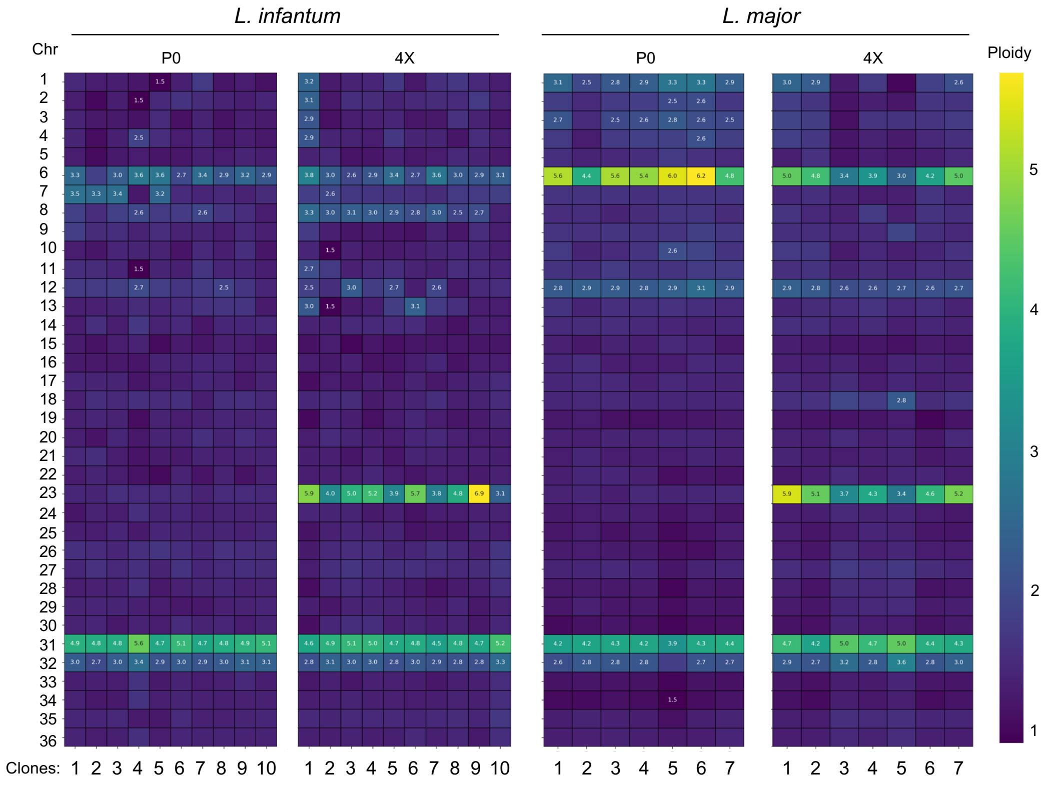


**Supplementary Figure 4.** Ploidy of *L. infantum* and *L. major* clones at P0 and 4X concentration.

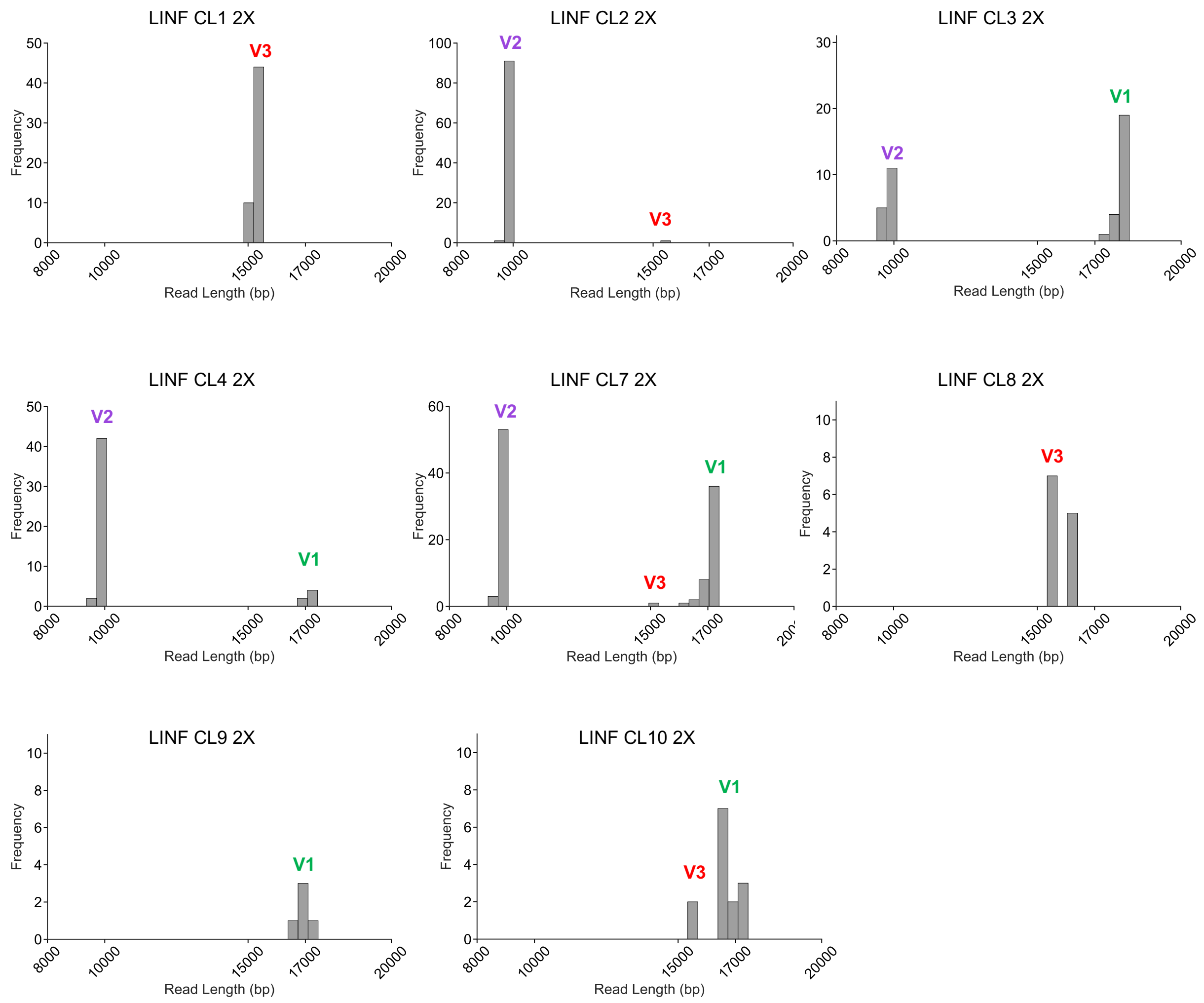


**Supplementary Figure 5.** Type 1 linear amplicon variants developed by LINF CL 1, 2, 3, 4, 7, 8, 9, and 10 at 2X concentration.

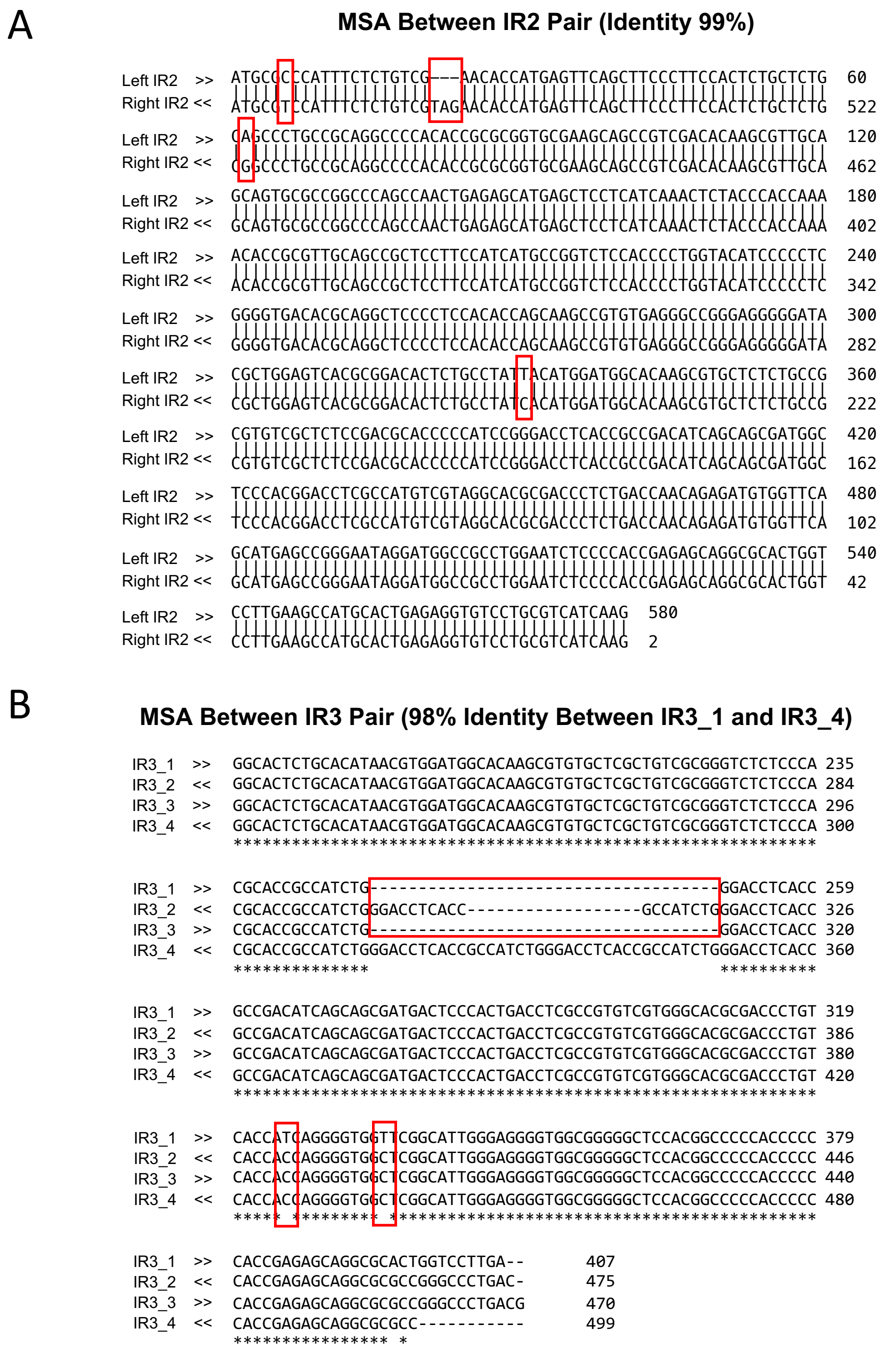


**Supplementary Figure 6.** The inverted repeat pairs, IR2 and IR3 are not 100% identical. **A)** Mismatches and InDels between the left and right IR2 (99% identical). **B)** There are four IR3s, two in the forward (IR3_1 from 160216 to 160623 bp and IR3_3 from 210095 to 210565 bp) and two in the reverse orientation (IR3_2 from 197650 to 198125 bp and IR3_4 from 212642 to 213141 bp). Among the four, IR3_1 and IR3_4 inverse repeat pairs performed recombination as reflected by the unique combination of SNPs and 36 bp InDel appearing on IR3_4 when ONT reads are mapped to the reference.

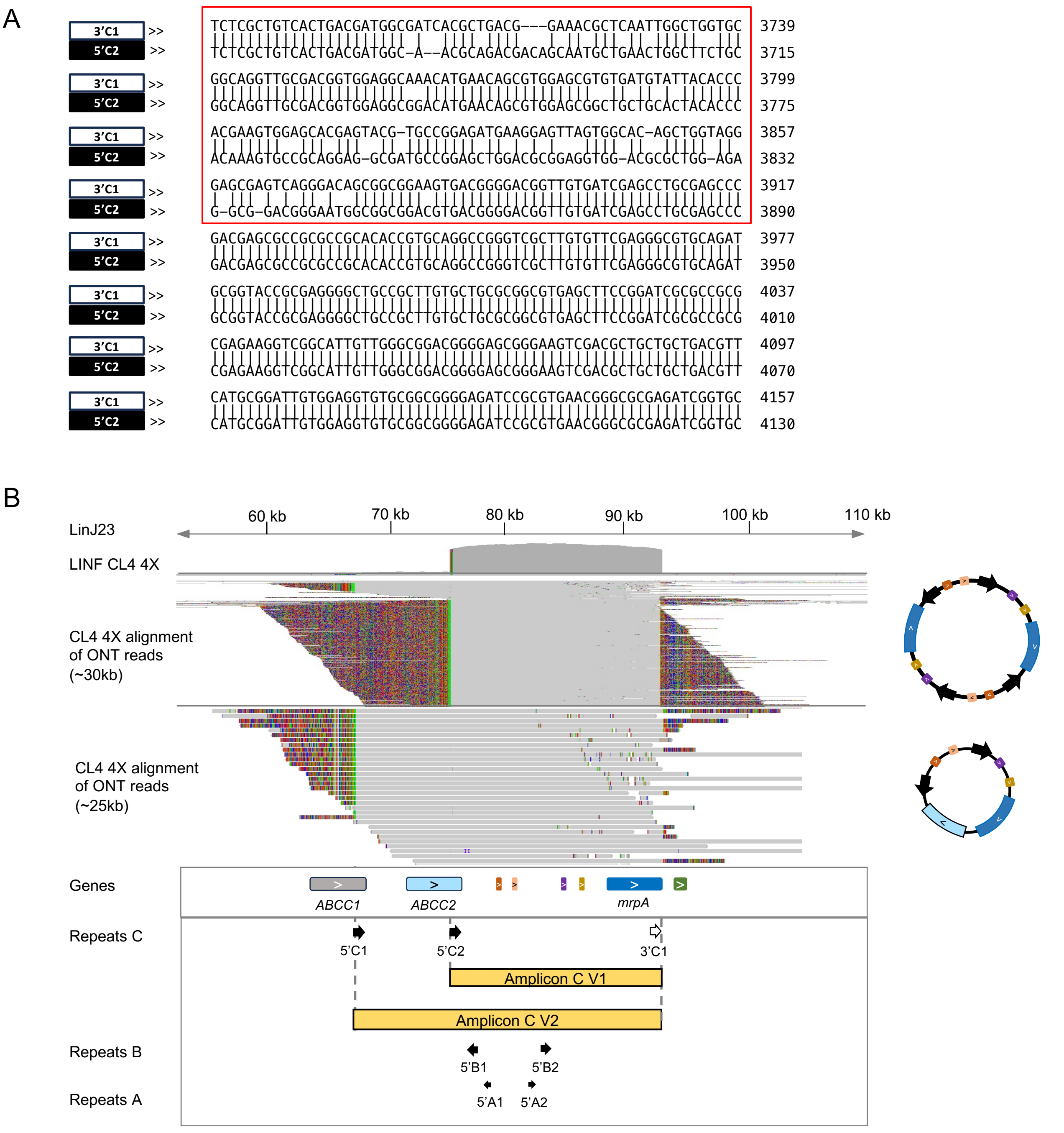


**Supplementary Figure 7. A)** Direct repeat C is present twice at the 5' side and once at the 3' side. The region in repeat C carrying InDels are marked in a red line box. **B)** Presence of the single copy amplicon C (~25 kb) along with the double amplicon C in LINF CL4 4X. 5'C1 and 3'C1 direct repeats participate in homologous recombination to form this amplicon.

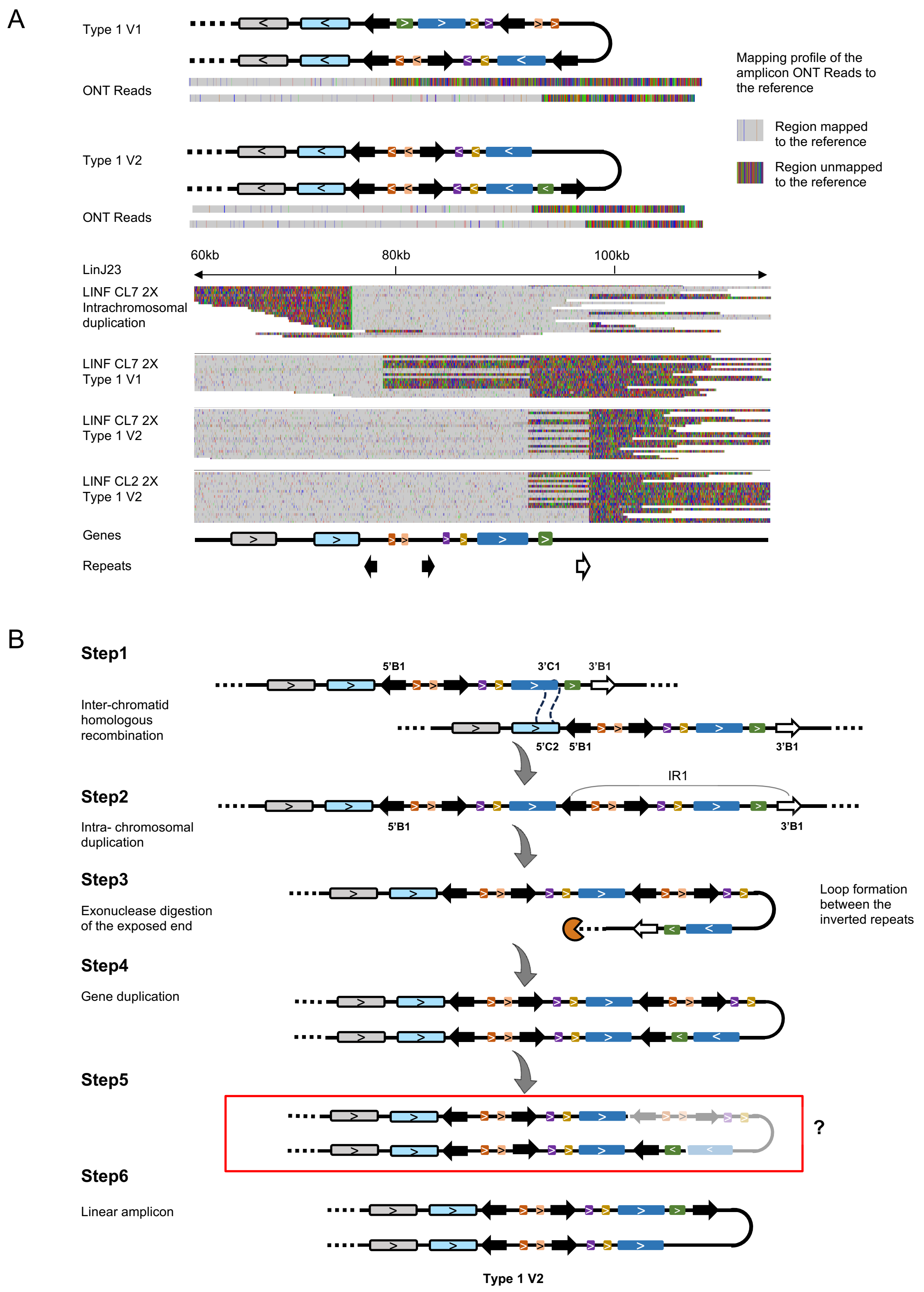


**Supplementary Figure 8. A)** Alignment of ONT reads from the intrachromosomal homologous duplication and two variants of type1 linear amplicons to the *mrpA* locus in CL7 2X. **B)** Mechanism of Type 1 V2 linear amplicon formation. Step 5 with red box is unconfirmed.

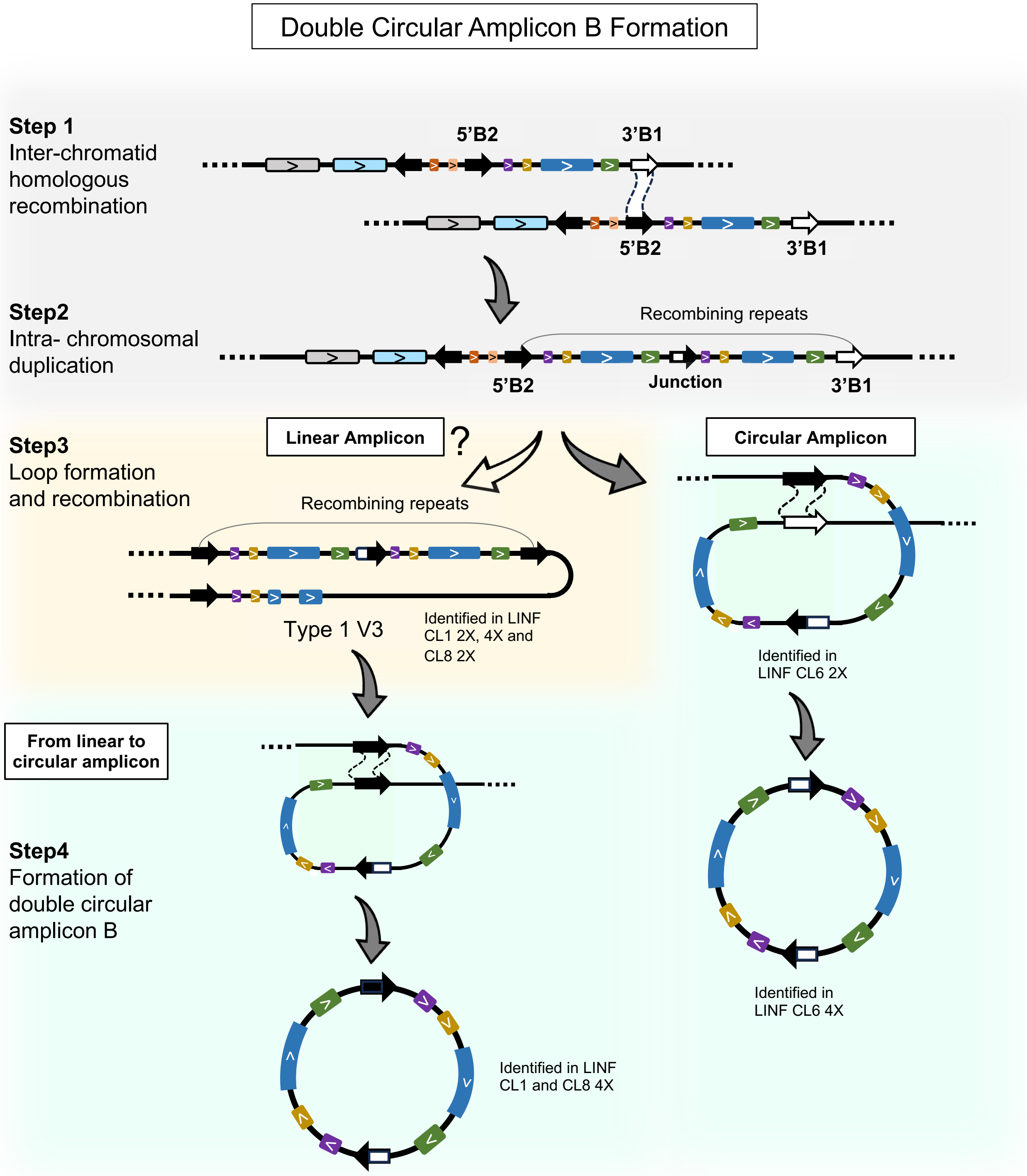


**Supplementary Figure 9.** Mechanism of double amplicon B formation in LINF CL1, 6, 8 at 4X. CL6 did not develop linear amplicon at 2X, but one read was detected with chromosomal duplication. CL1 and 8 developed linear amplicon Type1 V3 at 2X and double amplicon B at 4X indicating the progressive amplicon development from linear to circular with increasing Sb concentration. Development of linear amplicon Type1 V3 from chromosomal duplication could not be confirmed (indicated by the white arrow).

| *L. infantum* Clones | Linear Amplicon Type 1 V1 | Linear Amplicon Type 1 V2 | Linear Amplicon Type 1 V3 | Linear Amplicon Type 2 | Linear Amplicon Type 3 | Single Circular Amplicon A | Single Circular Amplicon B | Double Circular Amplicon B | Double Circular Amplicon C V1 | Single Circular Amplicon C V2 |
|---|---|---|---|---|---|---|---|---|---|---|
| CL1 P0 | | | | | | | | | | |
| CL1 1X | | | | | | | | | | |
| CL1 2X | | | | | | | | | | |
| CL1 4X | | | | | | | | | | |
| CL2 P0 | | | | | | | | | | |
| CL2 1X | | | | | | | | | | |
| CL2 2X | | | | | | | | | | |
| CL2 4X | | | | | | | | | | |
| CL3 P0 | | | | | | | | | | |
| CL3 1X | | | | | | | | | | |
| CL3 2X | | | | | | | | | | |
| CL3 4X | | | | | | | | | | |
| CL4 P0 | | | | | | | | | | |
| CL4 1X | | | | | | | | | | |
| CL4 2X | | | | | | | | | | |
| CL4 4X | | | | | | | | | | |
| CL5 P0 | | | | | | | | | | |
| CL5 1X | | | | | | | | | | |
| CL5 2X | | | | | | | | | | |
| CL5 4X | | | | | | | | | | |
| CL6 P0 | | | | | | | | | | |
| CL6 1X | | | | | | | | | | |
| CL6 2X | | | | | | | | | | |
| CL6 4X | | | | | | | | | | |
| CL7 P0 | | | | | | | | | | |
| CL7 1X | | | | | | | | | | |
| CL7 2X | | | | | | | | | | |
| CL7 4X | | | | | | | | | | |
| CL8 P0 | | | | | | | | | | |
| CL8 1X | | | | | | | | | | |
| CL8 2X | | | | | | | | | | |
| CL8 4X | | | | | | | | | | |
| CL9 P0 | | | | | | | | | | |
| CL9 1X | | | | | | | | | | |
| CL9 2X | | | | | | | | | | |
| CL9 4X | | | | | | | | | | |
| CL10 P0 | | | | | | | | | | |
| CL10 1X | | | | | | | | | | |
| CL10 2X | | | | | | | | | | |
| CL10 4X | | | | | | | | | | |
| Sb2000.1 | | | | | | | | | | |

**Supplementary Table 1.** Summary of amplicons formed in each LINF clone under four Sb drug concentrations. Each amplicon type is colored differently, where the darker shades of the color represent the dominant versions of the amplicons in case multiple versions of the same amplicon type are present in the same clone.
